# Mediterranean dispersal imposed bottlenecks in Neolithic sheep and goats

**DOI:** 10.64898/2026.07.21.739115

**Authors:** Áine Halpin, Jolijn A. M. Erven, Roger Alcàntara Fors, Conor Rossi, Andrew J. Hare, Valeria Mattiangeli, Simon Stoddart, Jarosław Wilczyński, Àngel Bosch Lloret, Raquel Piqué Huerta, Antoni Palomo Pérez, Josep Tarrús Galter, Xavier Terradas, Maria Saña-Seguí, Daniel G. Bradley, Kevin G. Daly

**Author notes:** contributed equally (Áine Halpin, Jolijn A. M. Erven, Daniel G. Bradley, Kevin G. Daly).

## Abstract

The Neolithic period in western Eurasia involved major dispersal events, including the translocation of livestock into Europe. These domestic animal translocations likely featured founder effects, fundamentally reshaping their genetic landscape. One well-known example is the Mediterranean dispersal, hypothesised to involve rapid maritime translocation with small founding herds, yet the genetic consequences for the livestock themselves remain poorly understood. To investigate this, we generated 12 palaeogenomes of sheep and goat from La Draga and Cova de l’Avellaner, and coanalyzed these with other Neolithic Mediterranean assemblages, including 4 additional novel genomes. We observe a significantly lower genetic diversity and higher runs of homozygosity (ROH) in Iberian sheep and goat populations compared to other ancient European and southwest Asian populations. We quantify the magnitude of the bottleneck in goats at ∼205 *Ne*, highlighting the severity of the founder event. We further identify species-specific demographic trajectories: goats experienced a severe and prolonged bottleneck, likely driven by regional isolation within the western Mediterranean, intensive exploitation strategies, and small initial founding herds, while sheep were likely more frequently exchanged across the region. Lastly, identity-by-descent (IBD) indicates connectivity along the Mediterranean coast compatible with a cabotage model. This suggests that seaways facilitated rapid initial dispersal but not sustained livestock transfer from southwest Asia, shaping the genetic foundation of Iberian livestock herds.

**Significance:** The Neolithic transition in western Eurasia fundamentally restructured human societies, yet the genetic consequences for translocated livestock remain poorly characterized. Analyzing 16 novel palaeogenomes from Mediterranean Neolithic sheep and goats, we demonstrate that maritime dispersal imposed severe founder effects on early Iberian livestock, reducing effective population sizes to as low as ∼205 in goats. Species-specific demographic trajectories reveal that goats experienced prolonged genetic isolation within the western Mediterranean, while sheep maintained broader regional connectivity. Identity-by-descent analyses support a cabotage model of coastal dispersal, indicating that Mediterranean seaways facilitated rapid initial colonization but not sustained genetic exchange with southwest Asian source populations. These findings reveal that the earliest livestock introductions into Europe were demographically constrained events whose genetic signatures persist in the foundations of Iberian pastoral heritage.

## Introduction

Domestic sheep and goats represent the first livestock to be brought under human control in southwest Asia (1–5), making them a key component of the agricultural practices which spread throughout Eurasia and Africa in the following millennia (3, 6). The onset of human control over these animals, combined with their repeated translocations, imposed demographic constraints that have shaped the genetic landscape of livestock populations. In particular, serial founder events associated with dispersal, linked to declines in genetic diversity, have been documented across many species (7–11). These include both goats (2, 12) and sheep (13), which exhibited high genetic diversity during early domestication (2, 12, 13). These dispersal bottlenecks arise through natural range expansions and more prominently human-mediated translocation (14–17).

One well-known dispersal process following the domestication of livestock in southwest Asia was their spread during the Neolithic period into Europe along two main migration routes: the Danubian and the Mediterranean (18). The expansion along the Western Mediterranean coast was rapid, with an estimated spread rate of 7.5-10.6 km/year (19, 20) corresponding to roughly 350 km per human generation (∼27 years) (19, 20). The speed of this dispersal is apparent in the earliest detections of Neolithic agricultural societies in different regions, reaching western Italy around 5,900–5,750 BCE (21, 22), southern France around 5,800–5,600 BCE (21, 22), and northeast Iberia around 5,700–5,600 BCE (23, 24). In contrast, the inland expansion into central Europe via the Danubian route progressed more slowly, with an average spread of approximately 1 km/year (25, 26). This difference highlights the distinctive nature of the Mediterranean dispersal, which likely depended on maritime mobility (27–29).

Due to the constraints of sea travel, the Mediterranean dispersal is theorized to have been small-scale (19, 20). Ancient human genome research on Early Neolithic Iberian and Maltese farmers has highlighted their low genetic diversity, which was hypothesised to reflect a bottleneck associated with the dispersal of farming populations along the Mediterranean coast (30, 31). Moreover, a reduction of domestic crop diversity was also observed along the Mediterranean dispersal route (27). However, little is known about how this process impacted livestock populations, which were presumably translocated in small numbers alongside migrant farmers, constrained by the capacity of their waterborne vessels and ability to sustain animals during voyages. A small number of palaeogenomic studies offer insight into this process, showing either high mitochondrial diversity in goats in Neolithic southern France (32) or low nuclear genetic diversity in sheep from Iberia and southern France (33, 34). These contrasting patterns are suggestive of species-specific bottlenecks during their initial Mediterranean dispersal, but individual studies have lacked larger comparative nuclear palaeogenome datasets to confirm the degree of diversity decline or indeed differing patterns between species.

The first point of contact for these Neolithic pioneers entering Iberia was Catalonia in northeast Iberia, with archaeological evidence suggesting initial *Impressa* influences around 5,700–5,600 BCE (23). The Early Cardial Neolithic followed shortly, beginning around 5,550 cal BCE, characterized by a stable territorial presence in coastal and pre-coastal areas alongside the utilization of caves (23). From 5,300 BCE onward, an increase in settlement intensity and territorial occupation is observed, marking the emergence of the Early Neolithic settlement of La Draga in the northeasternmost part of the region, which became a pioneering settlement in a previously sparsely populated region (23, 35–37). La Draga was an open-air lakeside dwelling with two distinct construction phases dating to 5,300–5,000 cal BCE and 5,000–4,700 cal BCE, respectively (38). The wooden constructions of the first settlement were contemporaneous according to the dendrochronological data (38). No evidence of preceding Mesolithic populations in the area has been documented (39), and the site features the full so-called Neolithic package (40–42). Nearby, the cave "Cova de l’Avellaner" was used as a funerary site from ∼5,250–4,740 cal BCE, containing 19 inhumations and a substantial assemblage of ceramic and faunal remains (43). The funerary activity at Cova de l’Avellaner is contemporaneous with the latest occupation phases of La Draga, and their close spatial proximity places them within the same Early Neolithic landscape of northeastern Iberia. Thus, palaeogenomes from La Draga and Cova de l’Avellaner would allow both the quantification of genetic consequences of the Mediterranean dispersal on its arrival to Iberia, and to assess our working hypothesis that the animals from the two assemblages derived from the same or related herds.

To investigate this, we generated palaeogenomes of sheep and goat from La Draga and Cova de l’Avellaner, in addition to other Neolithic Mediterranean assemblages to improve regional context (Table S1). Together, these genomes allow us to explore how the rapid Neolithic translocation impacted livestock species along the northern coast of the Mediterranean. Our results support the hypothesis that animals from La Draga and Cova de l’Avellaner share genetic signatures consistent with a shared herd, exchange networks, or broader regional demographic processes common across Early Neolithic herds in northeast Iberia. Additionally, we recover a demographic bottleneck in both species that aligns with the constraints imposed by maritime mobility during the Neolithic Mediterranean dispersal.

## Results

DNA was extracted from 12 sheep and goat remains from two Early Neolithic settlements in northeastern Iberia: Cova de l’Avellaner (5,250–4,740 cal BCE (43)) and La Draga (5,300–4,700 cal BCE (38)). We identified six sheep and two goats from Cova de l’Avellaner and one sheep and three goats from La Draga (Table S1). We additionally generated genome data from one goat from Sarakenos Cave (6,411-6,234 cal BCE) in Greece, two sheep from Le Taï (5,400–4,980 cal BCE (22)) in southern France, and one sheep from Santa Verna in Malta (3,600-900 BCE(44)). Following genome sequencing, final autosomal coverage ranged between 0.17-11.21✕ (median of 1.38✕). Molecular damage after uracil DNA–glycosylase (UDG) treatment and read length distributions were consistent with an ancient origin (Table S1). These new data (N=6 for goats, N=10 for sheep) were analysed with previously published ancient samples (N=99 for goats – Table S2, N=121 for sheep – Table S2), and modern data (Table S3-4 (45, 46)) We created a reference panel with published modern sheep, which was used for imputation, which was systematically evaluated (Supplementary Note S1, Figure S1-5). Additionally, some previously published samples were sequenced to a higher coverage (Table S2).

### Goats from La Draga and Cova de l’Avellaner share a strong genetic connection

To investigate the potential connection between La Draga and Cova de l’Avellaner (Figure 1), we evaluated multiple indicators of affinity between the recovered genomes. Placement of the Cova de l’Avellaner and La Draga goats on a principal component analysis (PCA) plot shows that they cluster closely together and are distinct from other Neolithic European genomes (Figure 2A). Relatedness analyses revealed no related individuals among these goats up to the method sensitivity threshold of third degree (Table S6 (47, 48)). A comparable but attenuated trend is evident in sheep, where the PCA placement (Figure 2A) of the Cova de l’Avellaner and La Draga genomes is shifted away from Neolithic southeast European and Anatolian sheep, forming their own distinct cluster. Mirroring the pattern seen in goats, no relatedness was found between sheep of these two settlements.

**Figure 1:**
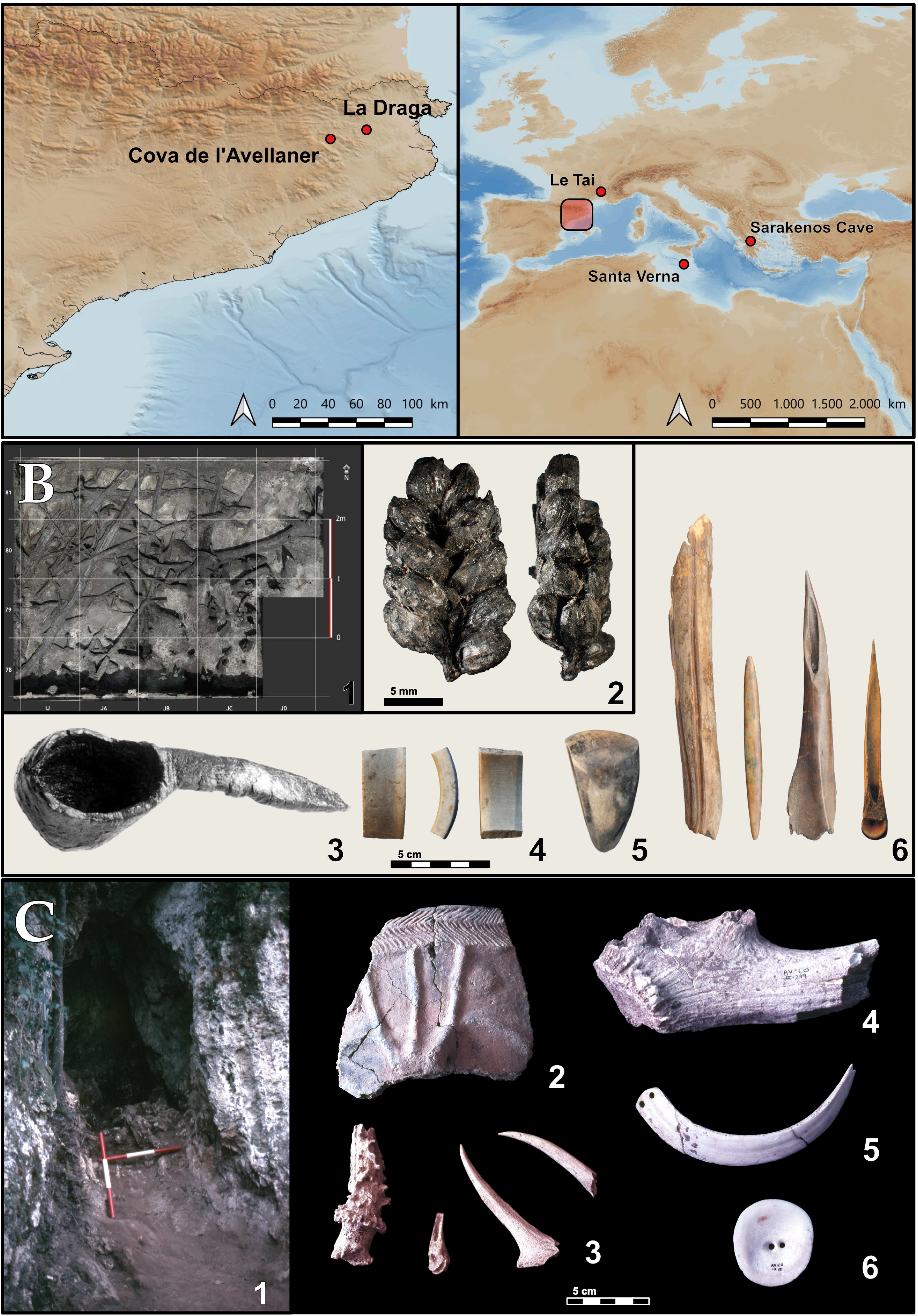
A) geographical map showing the location of La Draga and Cova de l’Avellaner in the northeastern Iberian Peninsula, and the other Mediterranean settlements. Base map from the IGN-CNIG WMTS service (MTN_Fondo layer; CC BY 4.0), derived in part from GEBCO Compilation Group (2021). Coastline data: LC 2023, Instituto Hidrográfico de la Marina (CC BY 4.0). Map prepared by the authors. B) La Draga. 1. Collapsed timbers from the wooden platform excavated in Sector D(54); 2. Durum wheat spike fragment; 3. Oak wood ladle; 4. Marble bracelet fragment; 5. Polished lithic adze blade; 6. Worked bone artefacts. Source: (2) Antolín, F.; (1-6) La Draga Archaeological Project. C) Cova de l’Avellaner. 1. View of Zone II; 2. Decorated fragment of Late Epicardial Vase 1 (Burial Cavity 1); 3. Roe deer antler fragment, a caprine metapodial awl fragment and roe deer antler awl fragments (Burial Cavity 1); 4. Fragment of deer antler with probable extraction for tool production (Burial Cavity 3); 5. Elongated biperforated wild boar tusk element (Burial Cavity 3); 6. Rounded biperforated wild boar tusk element (Burial Cavity 3). Source: Bosch, A. and Tarrús, J.

**Figure 2:**
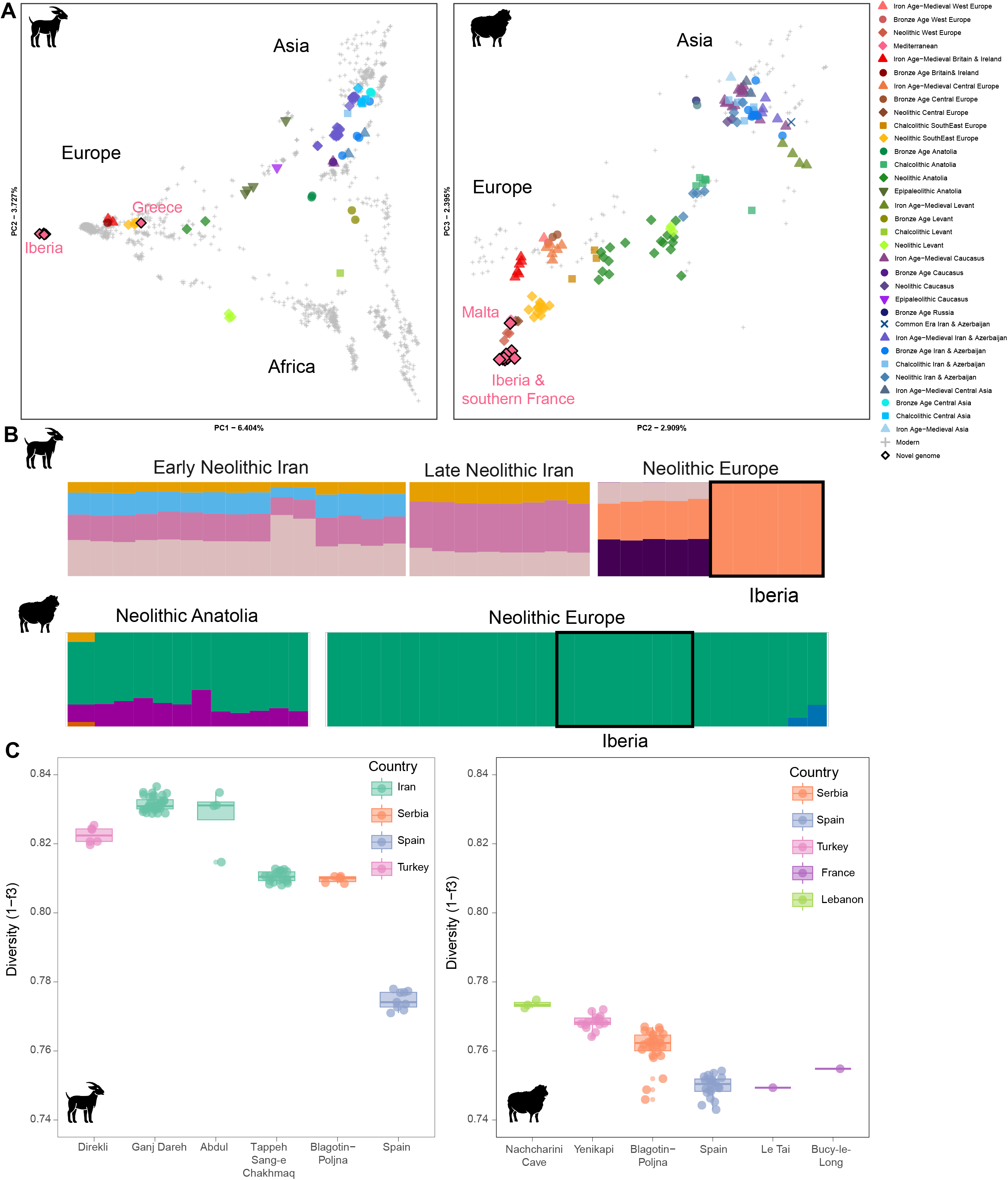
Population structure and genetic differentiation of goats and sheep. **A)** Principal Components Analysis using LASER, with ancient goats (N=75) and sheep (N=136) projected on modern variation. Key indicates geographical region and ancestry. **B)** ADMIXTURE profiles of Neolithic goats and sheep, all ancients are shown in Figures S6-7, and moderns are listed in Table S11-12 with an optimal K of 14 for goats and 6 for sheep. **C)** Within-population genetic diversities of ancient goats and sheep. Diversities were calculated using pairwise (1 − outgroup f_3_) between genomes from each site, with a minimum of 2 individuals per site.

We next quantified identity-by-descent (IBD) sharing, which can capture relatedness between individuals up to the eighth degree. Goats from La Draga and Cova de l’Avellaner showed a high level of IBD sharing, with all four (2 from each assemblage) exhibited relatedness above the 8th degree (Table S7). This is not reflected in sheep, where genomes from La Draga (N=1) and Cova de l’Avellaner (N=6) displayed IBD sharing but below this threshold (Table S8). The low degree of shared IBD suggests that the PCA clustering of these sheep genomes was more likely shaped by demographic history rather than close kinship. The difference in relatedness between sheep and goat implies species-specific management strategies or demographic backgrounds, with a smaller effective population size apparent in goats compared to sheep.

Altogether, these analyses corroborate a connection between La Draga and Cova de l’Avellaner: that goats from these sites likely originated from the same herd. The sheep show a more shallow genetic connection, potentially reflecting more extensive regional trade or animal exchange networks with neighbouring settlements, which is also consistent with a higher census count across Neolithic Iberia relative to goats (49–53). Consequently, the La Draga and Cova de l’Avellaner goats will be considered as one population hereafter, with sheep treated as distinct populations.

### Strong dispersal bottlenecks in Neolithic northeast Iberian sheep and goats

To evaluate the impact of Mediterranean dispersal on sheep and goat populations, we quantified genetic diversity and runs of homozygosity (ROH) across Neolithic assemblages. Calculating population diversity using pairwise combinations of (1 - Outgroup *f_3_*), we find the lowest diversity levels occurring within Neolithic Iberian populations in both species (Figure 2C, Table S9-10). In sheep, low diversity is also observed in the southern French settlement of Le Taï. The substantial decrease in genetic diversity within the Iberian and southern French populations hints at small local effective population sizes in both species (55–57). Ancestry modelling is also consistent with genetic decline affecting Neolithic Iberian herds (58, 59), with the Iberian genomes of both species assigned a single uniform ancestral component (Figure 2B, S6-7, Table S11-12).

In contrast, high genetic diversity is seen in early managed goats at Aceramic Neolithic populations in the Zagros Mountains (Ganj Dareh and Tepe Abdul Hosein), in mouflon at a hunted population from Nachcharini Cave in the PPNA (Lebanon, Figure 2), and in sheep at the Neolithic Turkish settlement of Yenikapı. Intermediate between these diversity extremes are Neolithic goats and sheep from Serbia (Blagotin-Poljna), which likely were translocated via mainland Europe rather than the Mediterranean.

Reduced genetic diversity in Neolithic Iberian livestock species was also reflected in an abundance of ROH, the genomic signatures of inbreeding (60). We found that the Iberian Cova de l’Avellaner and La Draga goats show high levels of small-to medium-sized ROHs (0.5-4 Mb, reaching on average 23.2% of the genome (14.2-30.8%)), while the number of large ROH (>4Mb) is relatively low (Figure 3, Table S12-13). These profiles insinuate a strong prolonged bottleneck, but without close-kin mating in recent generations (61–66). ROH totals of sheep from Cova de l’Avellaner and La Draga are amongst the highest observed, save for a single genome from Malta (Verna1), but also with little to no ROH tracts in the longest category (>16Mb), suggestive of a bottleneck rather than recent inbreeding (Figure 3).

**Figure 3.**
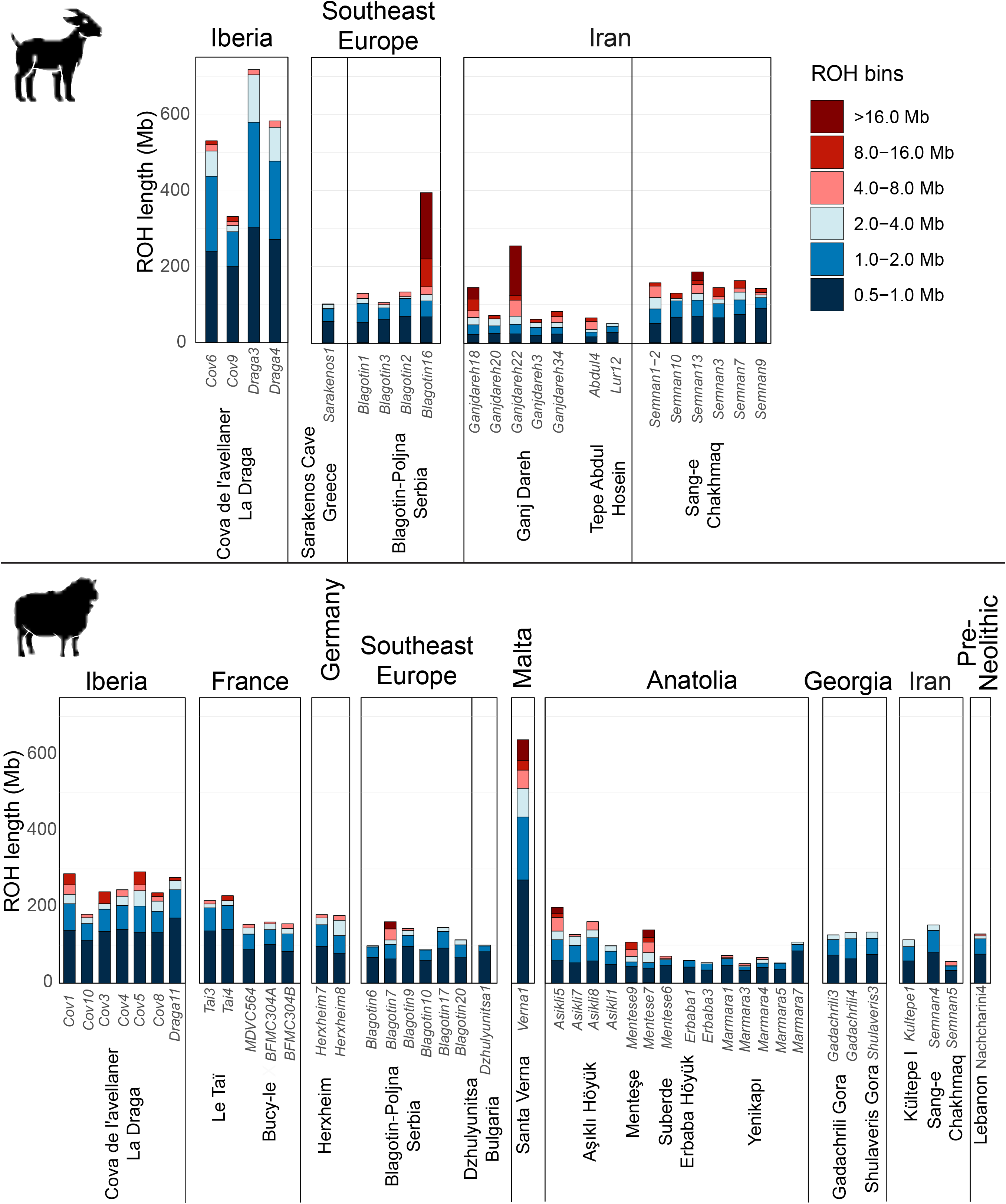
ROH profiles of goat and sheep. ROH length bin profiles for imputed ancient goats and sheep. ROH was calculated with each ancient sample individually, using transversions-only, mAF 5%, and downsampling to 1.2 million sites.

The absence of inbreeding signals may have been facilitated by herd management practices among the Early Neolithic communities associated with La Draga and Cova de l’Avellaner, which deliberately avoided mating between closely related animals. One possible strategy to limit inbreeding would be the control of sires: ethnographic research indicates that breeding males are often managed through controlled mating to reduce inbreeding and maintain diversity (67). Comparable ROH levels were seen within the French Le Taï sheep, suggesting shared dispersal history along the Mediterranean coast, consistent with cabotage (“To sail along the coast”). The observation that a published northern French sheep carries a lower amount of ROH than the Iberian and southern French goats may indicate it derived from distinct founding herds translocated via mainland Europe. This is consistent with broader patterns of ROH inferred here, where goats and sheep from central-southeast Europe (*p*=0.0159 and 0.0002, respectively) and southwest Asia (*p*=0.0004 and 2.45e-06, respectively) have significantly lower ROH compared to Iberia (Supplementary note S2).

Despite broadly similar trends, Iberian sheep and goats show marked differences in their absolute levels of ROH, where goats show extremely elevated profiles (61–66). A possible explanation is that relatively more sheep than goats were translocated via the Mediterranean route, or that the goat population of La Draga was small and isolated. At the Early Neolithic sites of Çatalhöyük and Aşıklı Höyük in Türkiye, sheep were likely more abundant (68–70), while in Neolithic Iberia, sheep also exceeded goats (49–53), suggesting that sheep outnumbered goats during the Mediterranean dispersal. However, the faunal spectrum of La Draga shows a roughly equal representation of sheep and goat (40, 71). A key difference between the species lies in their exploitation patterns: sheep exhibit mixed exploitation and longer survivorship curves, whereas goats were predominantly culled as young adults, suggesting a management strategy focused on meat production. Such management practices may have restricted the number of animals contributing to subsequent generations, particularly if only a limited number of males were used for breeding, thereby reducing the effective population size of the herd (72).

### Greater effective population size decline in goats than in sheep during the Neolithic Mediterranean dispersal

To assess the strength of the inferred bottleneck, we estimated effective population size in the Neolithic sheep and goats employing two complementary approaches: effective population size estimation using hapROH(73) and demographic simulations using SLiM5 (74). The ROH profiles inferred with hapROH mirror the patterns observed in our earlier analyses (Figure S8).The effective population size estimations for Iberian goats were 205 (95% CI 162-248, Std error 22), in contrast those from Serbia (Blagotin-Poljna) and the ancestral Iranian Zagros populations (Ganj Dareh) were estimated at 1,228 (95% CI 640-182, Std error 302) and 1,760 (95% CI 755-2,755, Std error 510, see Table S15), respectively. This corresponds to a 30% decline in effective population size from Zagros to Serbian goats, and an 83% decline between Serbian and Iberian goats. Simulation of inbreeding patterns in goats under domestication and dispersal conditions suggests a small population size leading to Iberian populations in the Neolithic (∼200-400 *Ne*, Supplementary Note S3, Figure S9-10), with population sizes dependent on population structure and genetic complexity during the initial dispersal from Asia. We also see a decline in effective population size in Iberian sheep (1386, CI=788-1,985, Std error 305) compared to Serbian sheep (5862, CI=714-11,050, Std error 2632) or those from the Early Neolithic Central Anatolian assemblage of Aşıklı Höyük (2,797, CI=715-4,872, Std error 1,061). However, the wide confidence intervals, reliance on a uniform recombination map, and a relatively small reference panel (N = 187), lead us to interpret these numbers with caution and refrain from between-species comparisons.

### Identity-by-descent analyses reveal the formation of the Mediterranean small ruminant gene pool

Identity-by-descent analyses provide the opportunity to quantify assemblage-level relatedness and more long-distance connections across the geographic range of the Neolithic dispersal. In general, sheep exhibit lower IBD sharing and relatedness than goats (Figure 4A). For example, at the Neolithic Serbian site of Blagotin-Poljna, relatedness is inferred in 30% of sheep, in comparison to 50% of goats. Similarly, within our Neolithic Iberian datasets, all goats are related to each other, while only 20% of sheep appear to be related. However, in both species, we see heterogeneity among individuals in their IBD sharing; on average goat share ∼235 cM in IBD, while sheep share on average ∼15 cM across the western Mediterranean (Figure S11-12, Table S7-8).

**Figure 4.**
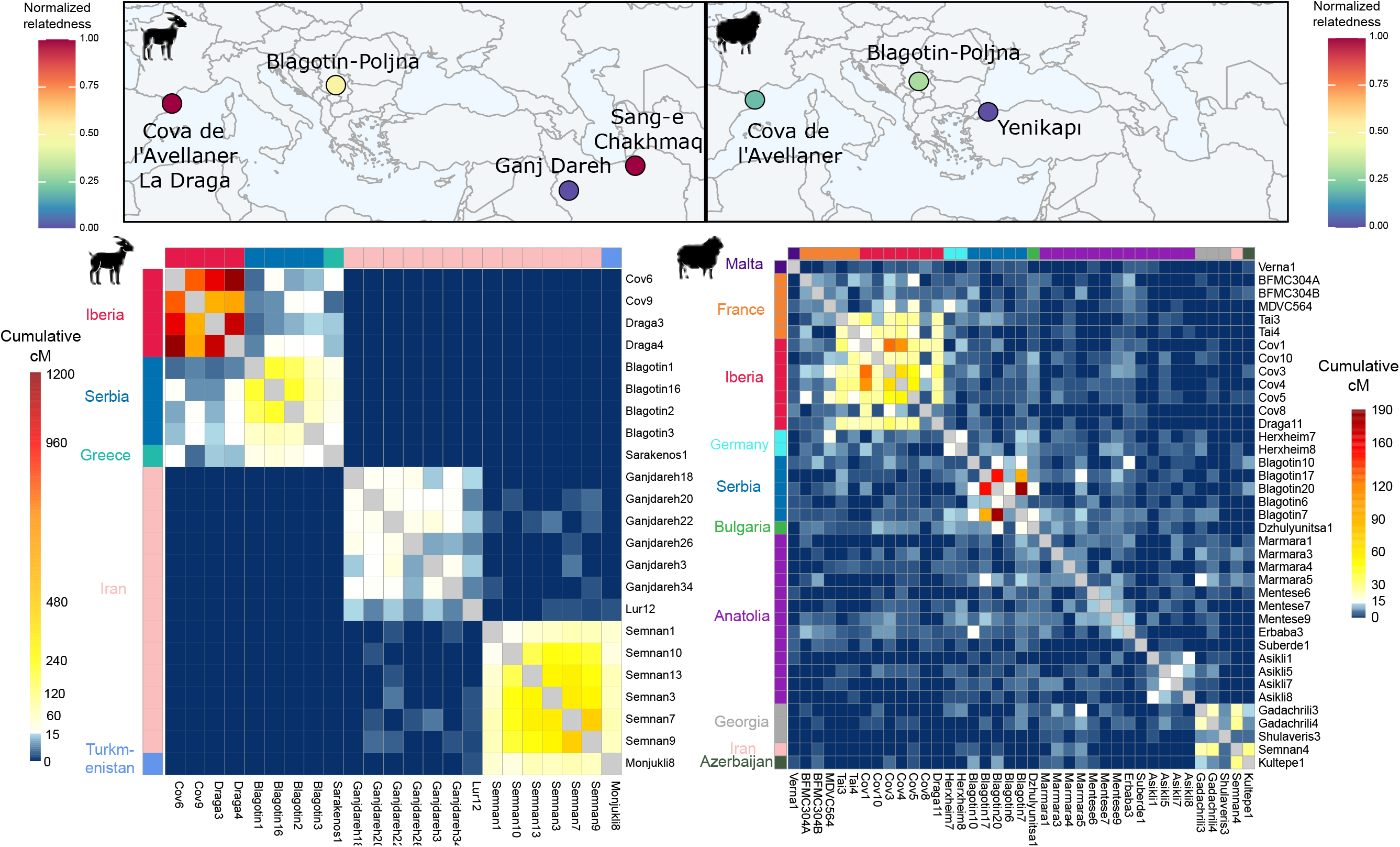
Identity-by-descent profiles for goats and sheep. **A)** Estimated for settlements with ≥3 samples, considering individuals related if they shared ≥3 segments >3 cM with a cumulative length of ≥21 cM. A normalized relatedness of 0 indicates no detectable genetic relatedness, whereas a value of 1 indicates that all individuals share genetic relatedness. **B)** Pairwise sharing of IBD, visualising IBD between individuals. Blue indicates little-to-no IBD sharing, and red indicates high IBD sharing; high IBD sharing clusters can be observed in Iberia.

We then assessed long-distance genomic connections by calculating between individuals >2cM IBD sharing across Neolithic Eurasia (Table S16-17). In goats, three distinct IBD clusters are observed (Figure 4), corresponding to within Europe, the Iranian Zagros Mountains, and northeast Iran/Turkmenistan. The Europe cluster is characterized by high IBD sharing between all Mediterranean goats from La Draga and Cova de l’Avellaner, which also share lower background IBD with goats from Serbia and Greece (Figure 4, S13). Sheep show a different pattern, with widespread background IBD sharing observed across the broader geographic range (Figure S14). We see only a single cluster of high IBD sharing within the western Mediterranean (Iberia and southern France, represented by the settlement of Le Taï), with the Neolithic northern French sheep (BFMC304A-B, MDVC564) not falling within this high IBD sharing cluster. On average, goats show higher IBD sharing in the western Mediterranean (∼870 cM in IBD) compared to sheep (∼26cM in IBD), reflecting species-specific differences in founding populations entering Europe. There is little-to-no IBD sharing between the Santa Verna sheep in Malta and other Mediterranean sheep: this is likely due to the time difference, with Verna1 being classified as Late Neolithic (or potentially Bronze Age, see Materials in the SI and Table S1). Elevated IBD sharing between northeast Iberia and southern France, a pattern that could reflect movement through Pyrenean corridors, is most convincingly explained by maritime connectivity along the coast.

## Conclusion

Our results provide evidence for a dispersal bottleneck which varied in its magnitude among species during the expansion of livestock populations along the northern Mediterranean coast in the Neolithic, which we quantify at around 205 *Ne*. This suggests that, although seaways may have acted as highways in the sense of rapid dispersal (19, 20), this did not facilitate sustained transfer of livestock from southwest Asia, thereby resulting in founder effects. The consequences of founder effects with limited gene flow is most apparent in the sheep recovered from Malta, whose long ROH tracts likely reflect an insular population with limited opportunity to intermix with other herds. This finding is mirrored in analysis of Neolithic humans from a nearby context (31). Despite the high level of ROH in the Santa Verna sheep, goats in Neolithic Iberia show even higher cumulative ROH levels, demonstrating the profound genetic impact of their translocation along the north Mediterranean coast. We further identify species-specific demographic trajectories: goats experienced a prolonged strong bottleneck, likely driven by regional isolation within the western Mediterranean, intensive exploitation strategies, and small initial founding herds. In contrast, sheep were likely more frequently exchanged within the region, with evidence of a maritime connection between northeastern Iberia and southern France. This pattern is compatible with a cabotage model (75), whereby migration took place via stepwise movements along successive coastal settlements. Neolithic maritime mobility was likely facilitated by dugout canoes, including well-preserved examples recovered from Italy (76, 77). Experimental archaeology has shown that such vessels could accommodate 9-11 people and were thought to have transported people, animals, and goods, although at relatively limited carrying capacities (76, 77).

## Methods

A full description of the materials and methods used are provided in the Supplementary Information appendix. Briefly, ancient DNA was extracted from 16 bone and tooth remains (6 goats and 11 sheep) in dedicated ancient DNA laboratory facilities at Trinity College Dublin. Samples underwent hypochlorite decontamination, EDTA predigestion, and overnight EDTA-proteinase K digestion, after which DNA was purified and used to construct double-stranded sequencing libraries (78). UDG treatment was performed prior to library construction (79). Libraries were sequenced on Illumina NovaSeq platforms at TrinSeq (Dublin) and Macrogen Europe (Amsterdam). Following adapter trimming and quality filtering with AdaptorRemoval version 2.3.2 (80), reads were aligned to the goat reference genome ARS1.0 (81) and the sheep reference genome ARS-UI_Ramb_v2.0 (82) using BWA aln version 0.7.17-r1188 (83). Reads with mapping quality below 30 and duplicate reads were removed. Species identity was confirmed using FastQ Screen (84), and coverage estimated with Qualimap2 (85).

Pseudohaploid genotypes were called by random read sampling using ANGSD (86), based on variant sites from the VarGoats reference panel for goats and an equivalent sheep reference panel (45, 87). Samples were additionally imputed using GLIMPSE2 (88) for goats and GLIMPSE1 (89) for sheep, applying a 100 kb-scale recombination map for goats (90) and a uniform recombination map for sheep (1cM=1Mb), and filtering for a genotype probability of 0.99. Final SNP datasets were restricted to transversions with a minor allele frequency of 5%, yielding 2,420,382 SNPs for goats and 1,302,568 for sheep. Molecular sex was determined from relative read depth across sex chromosomes, and pairwise relatedness assessed using PLINK (91) and READv2 (48).

Population structure was explored by projecting ancient individuals onto a PCA reference space built from modern populations using LASER v2 (92), with 100 replicates performed to ensure robust placement. Within-population genetic diversity was quantified using 1 − outgroup f₃ statistics computed with POPSTATS (93). Ancestry proportions in modern and ancient samples were inferred with ADMIXTURE v1.3.0 (Alexander et al., 2009), run across K = 2–16 clusters with the optimal K selected by cross-validation. We used a balanced dataset of goats (N=750), earlier described in *Lazzari et al. 2026* (46).

To assess inbreeding and effective population size, runs of homozygosity were computed from imputed genotypes using PLINK and bcftools (94), and independently with hapROH (73). Identity-by-descent segments were identified using RefinedIBD (95) prior by phasing with Beagle5(96), with segments merged using a sex-averaged recombination map and filtered for a minimum LOD score of 3 and length of 3 cM. Analyses were repeated three times to account for phasing variance.

To model the demographic impact of Mediterranean dispersal, forward-time population genetic simulations were performed in SLiM 5.0 (74) under a nucleotide-based whole-genome framework based on simHumanity (97), incorporating explicit recombination maps and goat demographic history. Bottleneck scenarios were simulated under panmictic and subdivided population models of varying complexity.

## Acknowledgements

The authors would like to thank Adamantios Sampson, Eve Rannamäe, Claire Manen, and Stéphanie Bréhard for providing access to the faunal remains and for information regarding the settlements. We thank the VarGoats sample providers acknowledged in *Denoyelle et al. 2021* and *Lazzari et al. 2026*. We also thank the metaAaRChive team for curating and publishing the metaAaRChive database, which we used to obtain metadata for ancient sheep and goat samples(98). This publication has emanated from research conducted with the financial support of Taighde Éireann – Research Ireland under Grant number 21/PATH-S/9515(T); and was supported by the European Research Council under the European Union’s Horizon 2020 research and innovation program (grant no. 885729-AncestralWeave). Archaeozoological analyses were supported by the ICREA Academia programme (MS) and the projects PID2020-115715GB-I00 and PID2023-152137NB-I00 (PI: MS), and contribute to the research activities of the EarlyFoods research group (SGR-Cat 2021, 00527).

## Data availability

The VarGoats genotypes are available at ENA under Project accession PRJEB90141. The sheep reference panel in vcf format will be deposited on zenodo and all ENA ids are in Table S5. The genetic data are deposited in the European Nucleotide Archive (ENA; accession no.PRJEB121319). Previously published data was used for this work.(2, 99–102).

