## Supplementary Methods and Figures S1-14 for "Mediterranean dispersal imposed bottlenecks in Neolithic sheep and goats"

### Material

#### *La Draga*

La Draga is an Early Neolithic lakeshore settlement located on the eastern shore of Lake Banyoles in northeastern Iberia (42.1267° N, 2.7586° E; 172 m a.s.l.), within a low-lying lacustrine environment (Palomo et al. 2014; Bosch et al. 2011). Radiocarbon dating indicates that the site was occupied between ca. 5,300 and 4,700 cal BCE. The earliest phase corresponds to the establishment of the settlement around 5,300 cal BCE and likely lasted only a few decades, followed by a hiatus and a later phase extending until ca. 5,000–4,700 cal BCE (Andreaki et al. 2022). Waterlogged conditions at the site have resulted in exceptional preservation of organic materials, including wooden structures and artefacts, plant remains, basketry, bone tools, lithics and Cardial pottery, providing detailed insights into the activities carried out within an early farming community (Terradas et al. 2017; Piqué et al. 2021; Antolín et al. 2014).

To date, 15,057 faunal remains have been analysed, of which 12,447 specimens could be taxonomically identified (NISP) (Ripoll Miralda 2025; Saña 2011). More than fifty animal species have been documented at the site (Ripoll Miralda 2025; Saña 2011), but domestic taxa dominate the assemblage, representing approximately 97% of the identified faunal remains (Saña 2011). The faunal record indicates a mixed livestock economy with a relatively balanced contribution of the main domestic taxa, including cattle (*Bos taurus*), pigs (*Sus domesticus*) and ovicaprines. Ovicaprines account for 43.6% of the identified remains, making them the most abundant domestic group with a sheep-to-goat ratio of ~1.0:0.8 and a minimum number of individuals of 27 sheep and 25 goats (Ripoll Miralda 2025). Sheep display a relatively broad mortality profile, suggesting a more polyvalent herd management regime, whereas goats are predominantly represented by young adult individuals, indicating a stronger focus on meat production. These patterns suggest that sheep and goats fulfilled complementary roles within a mixed livestock economy. Archaeobotanical evidence further documents the cultivation of crops such as barley and several wheat species, indicating that this livestock economy was closely integrated with agricultural production (Antolín et al. 2014).

#### *Cova de l’Avellaner*

The Cova de l’Avellaner is an Early Neolithic burial cave situated in northeastern Iberia (42°04′30″ N, 2°32′30″ E). Consisting of three natural apertures formed within a travertine outcrop, the site was adapted for use as burial spaces, separated by dry-stone walls (Bosch and Tarrús 1990; Gibaja et al. 2018). Radiocarbon dating suggests that funerary activity took place between c. 5,250 and c. 4,740 BCE, during the Early Neolithic (Gibaja et al. 2018). Previous genetic analyses of seven human individuals from the site revealed high mitochondrial haplotype diversity and a predominance of Y-chromosome haplogroup G2a among males, a lineage commonly associated with early Neolithic farming populations in Europe (Lacan et al. 2011). The archaeological assemblage includes pottery vessels, lithic artefacts, bone tools and personal ornaments made of shell, bone and stone, as well as faunal remains that are thought to be part of the funerary grave goods (Bosch and Tarrús 1990). The faunal assemblage comprises 1,769 identified specimens (NISP) and is dominated by domestic taxa. Ovicaprines represent the largest component (78.4%), followed by pigs (9.5%), dogs (3.75%), cattle (1.3%), red deer (5.6%), and roe deer (1.01%) with additional low frequency taxa (Molina i Serramitjana 1990). Within the ovicaprine sample, 927 remains (NISP), corresponding to 35 individuals (MNI), were identified and of the specimens attributable to species, 68.5% correspond to *Ovis aries* and 6.04% to *Capra hircus*. Most specimens lack cut marks or other anthropogenic modifications associated with butchery, suggesting limited or absent carcass processing prior to deposition. Age-at-death profiles based on epiphyseal fusion and dental wear stages indicate a predominance of e young individuals among the ovicaprines with only 16.7% reaching adulthood (Garcia-Reig 2016).

***Le TaÏ***

The Taï settlement (Remoulins) is located in the Gard departement, about 40 km from the Mediterranean Sea (43° 56' 34.19" N ; 4° 32' 34.87" E). The site is positioned in one of the numerous valleys deeply carved in the limestone plateau of Garrigues, opened to the plain of Remoulins. The site is at the interface of several ecosystems favourable to human life. It has been excavated for 10 years and thanks to the multidisciplinary research team it has been possible to understand sediment dynamics and general topography, to reconstruct the production system of the Neolithic communities and to discuss the functional status of each occupation (Manen 2022). The excavation took place in different sectors of the site: in the "cave", in its extension the "entrance" and in the open-air area in front of the cave. The history of the site, with its different phases has been investigated by field observations, the study of artefacts and 42 radiocarbon dates. The earliest human occupations belong to the Early Neolithic (EN) and the bayesian modelling shows that the first occupations of the site took place towards 5,400-4,980 cal BCE These layers are well preserved in the cave and in the "entrance" but have been totally eroded in the open-air sector. However, the dwellings probably extended to the open-air area and the cave could have been used for domestic activities and as a refuse area. The Early Neolithic layers (belonging to the Epicardial culture) have been divided in two ensembles (EN1 and EN2) according to the stratigraphy. The presence of numerous remains and structures argues in favour of a permanent settlement, and various activities have been identified: probable storage of cereals, cooking activities, knapping of flint and quartz, refuse areas, and so forth. Some ground stone tools indicate the presence of partially preserved zones of activity (food processing, sometimes in association with other undetermined activities) in the entrance of the cave and the porch. The use-wear analysis of the chipped flint industries demonstrates the practice of a wide range of activities dominated by craft processes on vegetal materials and hide but also involving hard animal and mineral materials and finally, harvesting and hunting activities perceptible through the maintenance of sickles and arrows. The zooarchaeological studies show a subsistence economy heavily based on ruminant herding and strongly suggest a self-sustaining system for the sheep, which were present on the site at least from the autumn to the end of the spring (Bréhard and Vigne 2022). The archaeobotanical remains indicate the cultivation of cereals but also the exploitation of wild plant resources from various plant communities around the site, particularly to be used as a food supplement for sheep/goats. These remains also suggest that the early Neolithic communities occupied the site in a permanent manner throughout the year.

***Sarakenos cave***

The Sarakenos Cave (38.4527°N, 23.2045°E) is an multicultural cave site located in the Kopais area in Boeotia, Greece (Sampson et al., 2009). Archeological records indicate that Sarakenos Cave was visited sporadically by human groups during the Final Paleolithic, but it changed during the Mesolithic and Early Neolithic when human visits were more frequent (Kaczanowska et al. 2016). Excavations at Sarakenos Cave unearthed several thousand animal bones - mainly remains of birds, medium size ruminants and rodents. Other groups of animals are less frequent - especially fishes or carnivores represented only by single bones. Faunal remains have been found in all layers, collected during the regular exploration, accompanied by not numerous archaeological materials. For our study whole bone material were separated on four main faunal assemblages: Early Neolithic (layer 2), Initial Neolithic (layer 3), Mesolithic (layer 4) and Palaeolithic/Pleistocene (layers 5-11). Systematic excavations in Sarakenos Cave have been conducted since the mid-1990s (Kaczanowska et al. 2016; Sampson 2008). The stratigraphic sequence was reconstructed in trench A where 12 layers were distinguished, Middle Paleolithic (12–11), Final Paleolithic (10–5), Mesolithic (4), Initial Neolithic (3), Early Neolithic (2), and younger deposits (Late Neolithic, Early Bronze Age). Layers 10–2 yielded several thousand animal bones.

During the excavations numerous and varied osteological materials were uncovered. The hard tissue remains of different taxa of molluscs, fishes, amphibians, reptiles, birds, and mammals were recovered in all layers, collected during the field work (Bocheński et al. 2018; Wilczyński et al. 2016). Early Neolithic layer 2 is dominated by presence of the goat/sheep bones and teeth, which are four times as frequent than the rest of animal remains, including birds bones. Among medium size ruminants, we could also mention single bones of domesticated pig and cattle (Wilczyński et al. 2016). In this layer remains of goats or sheep are very numerous (NISP = 236). They belong to a minimum of 4 individuals and represent all parts of the skeleton. Numerous are skull fragments, isolated teeth, vertebra and long bone fragments - especially hind limbs as well as phalanxes. The age structure of individuals included here may suggest that the material is dominated by juveniles killed at the age of >1.5 years old. Considering the animal remains, signs of human activity related mainly with consumption (burned bones created during roasting) are numerous, but also wholly burned bones were discovered. The season of death was nevertheless determined in three individuals discovered within layer 2 (Early Neolithic) showing a repeated pattern of late spring or summer kills (Wilczyński et al. 2016).

***Santa Verna***

The Santa Verna monument is located on the Xaghra plateau (36.0457 N, 14.2585 E; 141 asl) on the island of Gozo, Malta. The site was first excavated by Thomas Ashby in 1911, later by David Trump in 1961 and given a regional context by the Cambridge Gozo survey between 1987 and 1995 (Parkinson et al. 2020). The re-excavation as part of the FRAGSUS ERC project in 2015, directed by Caroline Malone, established conclusively that this was a so-called “temple” similar in size and structure to the more famous, and nearby, Ggantija monument. The stratigraphy of the monument runs throughout the Neolithic phases of the island, encompassing a probable gap between 4,800 and 3,800 BC. The main occupation of the site dates to the Ggantija period (c. 3,600 – 2,760 BC), but the sample was actually found in a probable destruction layer of the Bronze Age. For this reason, the sample has been directly dated in the Chrono Lab of Queen’s Belfast to: XXXXX

The petrous bone was chosen from a small sample of 348 individually identified bone fragments of sheep/goat (the dominant domesticate) across the site. The skull fragment was specifically found in Context 98, which contained two Neolithic ceramic snail figurines and many Neolithic sherds, but also four sherds of Bronze Age Borġ in-Nadur pottery (1675 -1225 BC). The context was located above Context (103) containing Tarxien (c. 2850-2660 BC) and Saflieni (c. 3080 -2760 BC) pottery and comprising a mid-brown compacted silty loam with mottled dark brown soil inclusions clearly derived from various clumps of loose deposit that had become mixed together through time (Parkinson et al. 2020).

### Methods

#### DNA extraction

DNA extraction was performed on all 21 bone remains (6 goats and 15 sheep) in dedicated ancient DNA laboratory facilities in Trinity College Dublin. Approximately 100 mg was obtained using a dental blade for each sample, cutting either the densest part of the bone element or from the available roots of each tooth. Specimens were then powderized using a MixerMill (Retsch).

DNA extraction followed a previously described pipeline (Mattiangeli et al. 2023). Briefly, samples were subject to a dilute sodium hypochlorite (0.5%) wash, followed by three $H_2O$ washes, and short EDTA predigestion (Damgaard et al. 2015), followed by an EDTA-proteinase K overnight digestion. The resulting supernatant was purified using High Pure Viral Nucleic Acid Large Volume purification kits Roche), eluting DNA in 50 µL of EBT (7.5 µL of 20% Tween in 15 mL EB buffer).

Purified DNA was then used to construct double-stranded DNA sequencing libraries (Meyer and Kircher 2010). Uracil bases were first excised (Briggs et al. 2010) by treating DNA with 5 µL Uracil-DNA glycosylase (UDG) over 3 h at 37 °C. DNA library construction then followed steps outlined in (Daly et al. 2018). In short, 3 µL purified DNA library was amplified using 1 µl of a unique P7 index oligo (5 µM) and 21 µL of amplification master mix (20.5 µL AccuPrime Pfx Polymerase (Invitrogen), 0.5 µL unique primer P5 (10 µM)). 12 cycles of amplification were performed. The amplified DNA was purified using Qiagen MinElute columns following manufacturer's instructions, eluting in 10 µL EB. Afterwards, amplified DNA was quantified using a Tapestation 4000 (Agilent). Additional DNA amplifications were performed at amplification cycles matching an expected 1 ng/µL concentration. The resulting amplifications were subject to high throughput sequencing on Illumina Novaseq 4000 (TrinSeq, Dublin) and NovaSeq X (Macrogen Europe, Amsterdam).

#### Data handling and genotyping

Adaptor trimming and read merging was performed using AdaptorRemoval version 2.3.2 (Schubert et al. 2016), removing collapsed reads shorter than 30bp and removing low quality terminal bases (--minadapteroverlap 1 --adapter1 AGATCGGAAGAGCACACGTCTGAACTCCAGTCAC --adapter2 AGATCGGAAGAGCGTCGTGTAGGGAAAGAGTGT --minlength 30 --trimns --trimqualities). Collapsed and raw read qualities were assessed using FastQC (Andrews and Others 2010). Specimen species were determined using FastQ Screen (Wingett and Andrews 2018), aligning reads against a set of genomes including sheep, goat, cattle, and human. We calculated coverage with Qualimap2 (Okonechnikov et al. 2016).

Following species ID (Table S1), reads from UDG-treated libraries of goat samples were aligned to the goat reference genome ARS1.0 (Bickhart et al. 2017), and sheep samples were aligned to the sheep reference genome ARS-UI_Ramb_v2.0 (Davenport et al. 2022). Using a pipeline previously described (Daly et al. 2021). Briefly, collapsed reads were aligned using bwa aln version 0.7.17-r1188 (Li and Durbin 2009) relaxing alignment parameters (-n 0.01 -o 2) and converted to bam files using samtools version 1.13 (Li et al. 2009). Using samtools, bam files were filtered to remove reads with mapping quality <30, and for duplicate reads.

Pseudohaploid genotypes for goat variant sites discovered in the VarGoats genome dataset (Denoyelle et al. 2021; Erven et al. 2025) were called by random read sampling using ANGSD (Korneliussen et al. 2014), previously described (Erven et al. 2025). Additionally, samples were imputed using GLIMPSE2 (Rubinacci et al. 2023) using a pipeline previously described (Erven et al. 2025), employing the VarGoats dataset as a reference panel and a 100 kb-scale recombination map as the genetic map (Etourneau et al. 2025). Imputed samples were filtered for a genotype probability (GP) of 0.99. Datasets were then further filtered to only contain transversions and a MAF of 5% for goats, resulting in 2,420,382 SNPs.

Pseudohaploid genotypes for sheep variant sites discovered in the sheep reference panel (See SI note 1: Imputation) were called by random read sampling using ANGSD, previously described (Erven et al. 2025). Additionally, samples were imputed using GLIMPSE1 (Rubinacci et al. 2021), SI note 1: Imputation. Datasets were then further filtered to only contain transversions and a MAF of 5% , resulting in 1,302,568 SNPs.

**LASER**

A projection Principal Component Analysis (PCA) using LASER v2 (Wang et al. 2015) was performed. The PCA reference space and projection transformation for goats were constructed using the VarGoats dataset, filtered to retain transversions with a MAF of 5%, resulting in a total of 2,420,382 SNPs. All ancient samples were then projected onto the PCA space, and then filtered for individuals covered by less than 10000 loci. To reduce stochastic variation and provide robust estimates, 100 replicates were conducted.

For sheep, a projection PCA was generated in a similar fashion. The PCA reference space and projection transformation was constructed using the sheep reference panel (SI note 1: Imputation). The reference panel was filtered to retain only transversions with a MAF of 5%. This resulted in a total of 1,302,568 SNPs. All ancient samples were then projected onto the PCA space, and then filtered for individuals covered by less than 5000 loci. To reduce stochastic variation and provide robust estimates, 100 replicates were conducted.

**1-Outgroup f3**

We estimated within-population genetic diversity using 1 − outgroup $f_3$ values based on the ascertained pseudohaploid SNP panels described above, restricting analyses to transversions, for both sheep and goat. Outgroup $f_3$ statistics (Patterson et al. 2012) were calculated using the Python program POPSTATS (Skoglund et al. 2015). Analyses were performed with the options *--f3vanilla* (computing $f_3 = (p_3 - p_1)(p_3 - p_2)$) and *--not23* (allowing the use of non-human chromosomes).

For each test, individuals were specified as *Individual1, Individual2, outgroup, outgroup*, with sheep used as the outgroup for goats and goat for the outgroup of sheep. Genetic diversity was then quantified as $1 - f_3$, following (Kaptan et al. 2024).

**ADMIXTURE**

Ancestry proportions in modern and ancient samples were inferred using ADMIXTURE v1.3.0 (Alexander et al., 2009). For goats, analyses were performed on the combined VarGoats and ancient dataset, and for sheep on the combined modern and ancient dataset described above. ADMIXTURE was run for values of K ranging from 2 to 16, and the optimal number of clusters was determined based on cross-validation (CV) error (--cv=10). The best K for goats was 14 and 6 for sheep.

**Runs-of-homozygosity (ROH)**

ROH were computed on goats and sheep using imputed genotypes using a combined plink and bcftools pipeline, previously described and validated (Erven et al. 2025). In summary, samples were downsampled to 1.2 million SNPs, filtered for minor allele frequency >5% and restricted to transversions. ROH were calculated for each individual using both PLINK v1.90 (Chang et al. 2015) and bcftools v1.17 (Danecek et al. 2021). Plink was run with standard parameters aside from setting the --homozyg-window-snp to 200. Bcftools was run with standard parameters aside from -G 30 --AF-dflt 0.4. Bcftools output was further filtered to retain regions containing at least 200 SNPs, a quality score greater than 10, and a minimum length of 500 kb. Long ROHs identified by bcftools (>4 Mb) were merged with the plink ROH profiles using mergeBed (Quinlan and Hall 2010), counting the number and keeping the sizes of merged ROHs with the parameters -c and -o count, collapse.

For hapROH the pipeline from (Ringbauer et al. 2021) was followed. First we converted the reference panel VCF into hdf5 format using h5py and scikit-allele in a custom python script. The recombination map was also added as information to the hdf5 file. Then we followed the hapROH pipeline described in the hapROH vignette (https://haproh.readthedocs.io/en/latest/tutorial.html). In short, we transformed plink files to eigenstrat with *convertf*. From the eigenstrat files, ROHs were calculated for each individual separately using the *hapsb_ind* function, adjusting the number of reference individuals (n_ref) and the conPop parameter. These individual files were then combined to a single ROH output file with the *pp_individual_roh* function. For the estimation of effective population size (Ne) we excluded inbred samples ( ≥50 cM in ROH >20 cM). The updated IDs and combined ROH file were then used as input to the *load_roh_vec* function. Both methods were tested, using the default parameters aside from the chromosome sizes (Goat and sheep chromosome sizes were used respectively).

**Identity-by-descent (IBD)**

Identity-by-descent segments were identified using a similar pipeline as described in (Erven et al. 2025). In summary, imputed genotypes were phased together using Beagle5 (Browning and Browning 2013) with standard parameters aside from setting impute = true and specifying a random seed. Genotypes below GP99 were set to missing, and SNPs were further filtered for the 1,037,536 SNPs used in Erven et al. (2025) (MAF 5%, transversions and had no missing data in imputed samples >2×), and finally filtered for no missingness.

RefinedIBD (Browning and Browning 2013) was run with default settings. To correct for breaks or short gaps in IBD segments, segments were merged with a sex-averaged 100 kb-scale recombination map (Etourneau et al. 2025), using merge-ibd-segments.17Jan20.102.jar, allowing a maximum of one discordant homozygote and gaps less than 0.6 cM and 4cM in length. The phasing, refinedIBD and merge-ibd were repeated a total of 3 times to account for potential variance in phasing. The first 2 Mb of chromosome 18 was excluded as in Erven et al. (2025). The repeated runs were combined and filtered for a minimum LOD score of 3 and a length threshold of 3 cM. To obtain IBD segments for a pair of individuals, IBD segments for pairs of individuals were extracted from the IBD files and merged with bedtools MergeBed.

A similar approach was taken for sheep, imputed genotypes were phased together using Beagle5 with standard parameters aside from setting impute = true. The SNPs were then filtered with a MAF filter of 5%, transversions only and filtered for no missingness. RefinedIBD, segment merging, repeated phasing runs, and downstream filtering were then carried out as described above.

**Forward simulations with SLiM**

We performed forward-time population genetic simulations using SLiM 5.0 under a nucleotide-based, whole-genome framework. The simulation followed a SimHumanity-style approach as described by Haller et al. (2025), enabling explicit modelling of mutations, recombination, and demography. Tree-sequence recording was enabled with periodic simplification to efficiently store the ancestral recombination graph.

The genome was modelled at nucleotide resolution using a Jukes–Cantor mutation model and a sex-averaged 100 kb-scale recombination map (Etourneau et al. 2025). Chromosome (chr28) was simulated with explicit genomic coordinates and a randomly initialized ancestral sequence.

The simulation was initialized with a diploid ancestral population (N = 10,000), followed by burn-in and subsequent demographic events including population size changes, splits, and bottlenecks following goat demographic history. At defined time points, the Mediterranean bottleneck was simulated, this was done for a single panmictic population, 3 subpopulations, and 6 subpopulations. All scripts are made available at https://github.com/JolijnErven.

The tree-sequence output was converted into a VCF file using the Python packages tskit, msprime, and pyslim. First, the tree sequence was loaded using *tskit.load*, alleles were then reconstructed using *pyslim.convert*, and finally the VCF file was generated using *write_vcf*. The subsequent VCF was used for ROH reconstruction.

**Mitochondrial genomes**

Reads were aligned to a circularised version (Daly et al. 2018) of the goat mtDNA reference genome (NC_005044.2; 15bp of each side concatenated to the opposite end) using parameters described under "Data Handling and Genotyping". mtDNA sequences were called using ANGSD version 0.937-74-g9100f3d (Korneliussen et al. 2014), using -doFasta 2 with the following parameters: minQ 20 -minMapQ 30 -trim 4 -setMinDepth 3. For samples with <4× mtDNA coverage, -setMinDepth 1 was used instead. The resulting fasta files were decircularized using a python script by removing the first and last 15bp of the sequence. mtDNA sequences were aligned with published mtDNA sequences using MUSCLE (Edgar 2004) and ML phylogenies constructed using phyML (Guindon et al. 2010) through Seaview

(Gouy et al. 2010). After initial phylogenetic placement, samples were realigned to a closer haplogroup sequence (Daly et al., 2018) and final mtDNA sequences generated as above.

**Molecular sex**
Molecular sex was determined using the relationship between the number of aligning reads to each chromosome vs. chromosome length (Park et al. 2015). Male samples compared to female samples are expected to show lower read alignment to the X chromosome relative to its length, due to the single copy of the X chromosome expected to be carried. This was performed the same for sheep and goats.

**Relatedness analyses**
Pairwise genetic relatedness among individuals was assessed using two complementary approaches. With --genome function implemented in PLINK and READv2 (Alaçamlı et al. 2024), estimating kinship categories from pairwise allele sharing. Both methods were performed on the pseudohaplotypes for both sheep and goat and results were compared to identify and validate close biological relationships within the dataset.

### SI note 1: Building sheep reference panel and systematic evaluation of sheep imputation

#### Modern Genome Processing and reference panel compilation
Publically available fastq files for 200 sheep from diverse geographic origins  (Table S4) were downloaded through the ENA browser. FastQC was carried out on each file to confirm the absence of adapter sequences and for general quality control (Andrews and Others 2010). Reads were aligned to ARS-UI_Ramb_v2.0 (Davenport et al. 2022) using BWA mem ver.07.13 (Li and Durbin 2009). Reads were sorted and quality filtered (-q 30) with SAMtools ver. 1.9 and duplicate reads were removed using Picard ver 2.20.3.

Variants were called from the 200 aligned modern sheep using Graphtyper (Eggertsson et al. 2017a). Graphtyper was run per chromosome. The variants were then subjected to filtering,  largely following the parameters that are detailed in (Eggertsson et al. 2017b; Erven et al. 2024). Using VCFTOOLS ver.0.1.17 (Danecek et al. 2011), SNPs were removed if they were within 3bp of another variant, INDELs were removed also. SNPs were further filtered for biallelic alleles, a minimum depth of 6x, and a maximum genotype depth of 3x the average genomic coverage of the panel. Vcftools was also used to apply a minimum quality filter of 25 as well as a genotype quality filter of 20. The bcftools filter was used to filter the panel according to Graphtypers guidelines with (QD > 2., SB < 0.8, MQ > 40.0, LOGF > 0.5, AAScore > 0.5; (Li 2011). Vcftools was finally used to filter out singletons, repetitive regions and genotypes with greater than 20% missingness. Following this, the individuals in the panel were analysed for missingness and relatedness with other panel members. Any individuals with more than 20% missingness were removed. Any individuals with third degree or higher relatives were also removed from the panel.

This filtering resulted in the panel containing 187 individuals and 25,649,678 million SNPs. The reference panel was then phased using Beagle with parameters *impute=false, window=40, overlap=4, gp=true ne=20000*.

**Imputation**
Two ancient sheep and two modern genomes were downsampled to 0.3, 0.5, 0.7 and 1.0x per chromosome. The chromosomal coverages were estimated using Qualimap2 (Okonechnikov et al. 2016). Picard ver 2.20.3 was used to downsample each chromosome to the desired genomic coverage.

The GLIMPSE version 1.1.1 (Rubinacci et al. 2021) pipeline was used to generate genotype calls and likelihoods with bcftools mpileup (version 1.12) with parameters -I, -E, -a "FORMAT/DP,FORMAT/AD,INFO/AD", the reference genome and the SNPs in the reference panel (-T) followed by bcftools call with the parameters -Aim -C alleles, and the reference panel sites (-T). This step was performed on both downsampled and high-coverage genomes.

The downsampled genomes were imputed using the GLIMPSE pipeline. Chromosomes were split into chunks of 2 Mb with a 200 kb buffer window with GLIMPSE_chunk. Imputation was performed on these chunks with default parameters using GLIMPSE_phase with the reference panel created in the section variant calling. The imputed chunks were ligated using GLIMPSE_ligate with default parameters. Following ligation of chunks, chromosome files were merged. With bcftools, the data was filtered for GP≥0.99 and INFO ≥0.99. The high-coverage genotype likelihoods were further filtered for a minimum base quality of 30, a minimum genotype quality of 25, a minimum genotype coverage of 7x , a maximum genotype coverage of 3 times the average genomic coverage, and a minimum allelic balance of 40%.

**Concordance and accuracy**
To investigate genotype accuracy, concordance between high coverage genotypes and their downsampled counterparts was measured. This was done using an in-house python script which computes concordance between two VCF files; a high quality vcf and a downsampled imputed vcf. It measures correctly imputed, incorrectly imputed and concordance and site recovery rates. Accuracy was investigated for both homozygous alternative (i.e non-reference) and heterozygous sites. Accuracy was investigated across various MAF bins and at a 2.5 MAF threshold. (SI fig. 1) Concordance values were verified using bcftools gtcheck. Error rates were also calculated by dividing incorrectly imputed by total genotypes.

The proportion of missingness was also investigated across the downsampled individuals. This was calculated by 1-(present SNPs/total sites). (SI fig. 2-3)

**Runs of Homozygosity (ROH)**
To test consistency of imputed genomes in analysis, ROH were estimated across both the high-coverage genome and its downsampled imputed counterparts. (figure S4) This analysis was carried out using PLINK v1.90 with parameters--homozyg --homozyg-density 50 --homozyg-gap 100 --homozyg-kb 500 --homozyg-snp 50 --homozyg-window-het 1 --homozyg-window-snp 50 --homozyg-window-threshold 0.05 --sheep. (Chang et al. 2015)

**Principal component analysis (PCA)**
Principal component analysis (PCA) was performed to further assess the genetic similarity between high-coverage and imputed genotype datasets. (figure S5) Imputed datasets were filtered to retain only high-confidence genotype calls with genotype probability (GP) ≥ 0.99.

Variants with a minor allele frequency (MAF) below 2.5% were excluded prior to analysis. The filtered variants were converted to PLINK format and merged into a single dataset. Linkage disequilibrium (LD) pruning was performed using PLINK v1.9 with the indep-pairwise command (window size = 50 SNPs, step size = 5 SNPs, $r^2$ threshold = 0.2), and PCA was calculated from the resulting LD-pruned marker set. Principal component coordinates were obtained using the PLINK --pca function and visualized in R (v4.x) using the ggplot2 package. Samples were coloured according to individual and sequencing/imputation status to facilitate comparison between high-coverage and imputed genotypes.

**SI note 2: Statistical analysis of ROH profiles**

A goat from Sarakenos Cave in Greece (6,411–6,234 cal BCE), representing a population that did not go through the dispersal bottleneck along the Mediterranean coast, lacks the high number of short-to-medium ROH seen in Iberian goats (Sarakenos 102 Mb, and Iberia on average 520 Mb). This trend is similar for other Neolithic southwest Asian and European goats, which shows a significant lower ROH profile compared to Iberia ( Wilcoxon rank sum exact test for Southeast Europe, southwest Asia: W=19 and 52, one-tailed p=0.01587 and 0.0004202, respectfully). A similar pattern is observed for Neolithic sheep, where sheep from central and eastern Europe (Wilcoxon rank sum exact test; W=54, one-tailed p=0.0001998), Anatolia (Wilcoxon rank sum exact test; W=125, one-tailed p=2.447e-06), and other parts of southwest Asia (Wilcoxon rank sum exact test; W=54, one-tailed p=0.0001998), show a significantly lower ROH signal than Iberia.

**SI note 3: Modelling demographic scenarios in goat using SliM5**

We modelled potential demographic histories using chromosome 28 in goats, first incorporating a bottleneck consistent with hapROH results (Ne of 200). The overarching demographic model followed SimHumanity (Haller et al. 2025), with parameters adapted to goat demographic history. A bottleneck with an Ne of 200 is inconsistent with our ROH profiles, as simulated individuals show elevated ROH and reduced heterozygosity; this stabilizes with increasing Ne, and simulations match empirical ROH profiles at approximately Ne ~500 (Figure S9). This single bottleneck scenario may be overly simplistic, so we simulated migration with population structure toward the western Mediterranean, instead of a single homogeneous founder population. This scenario matches the empirical data better with an overall Ne of ~400 for 3 subpopulations (133 Ne per subpopulation) and an Ne of ~250 for 6 subpopulations (~42 Ne per subpopulation; Figure S10). This illustrates the effect that population structure (if herds were drawn from multiple distinct genetic units in Anatolia rather than a single homogenous group) would have on the ROH profiles of European founder flocks. However, this represents only an exploration of some possible, and more in-depth studies could reveal the full complexity of the Mediterranean migration. Overall, the effective population size of goats during the Mediterranean migration was low (~200-500 Ne), and livestock were likely translocated with multiple flocks.

### Supplementary figures

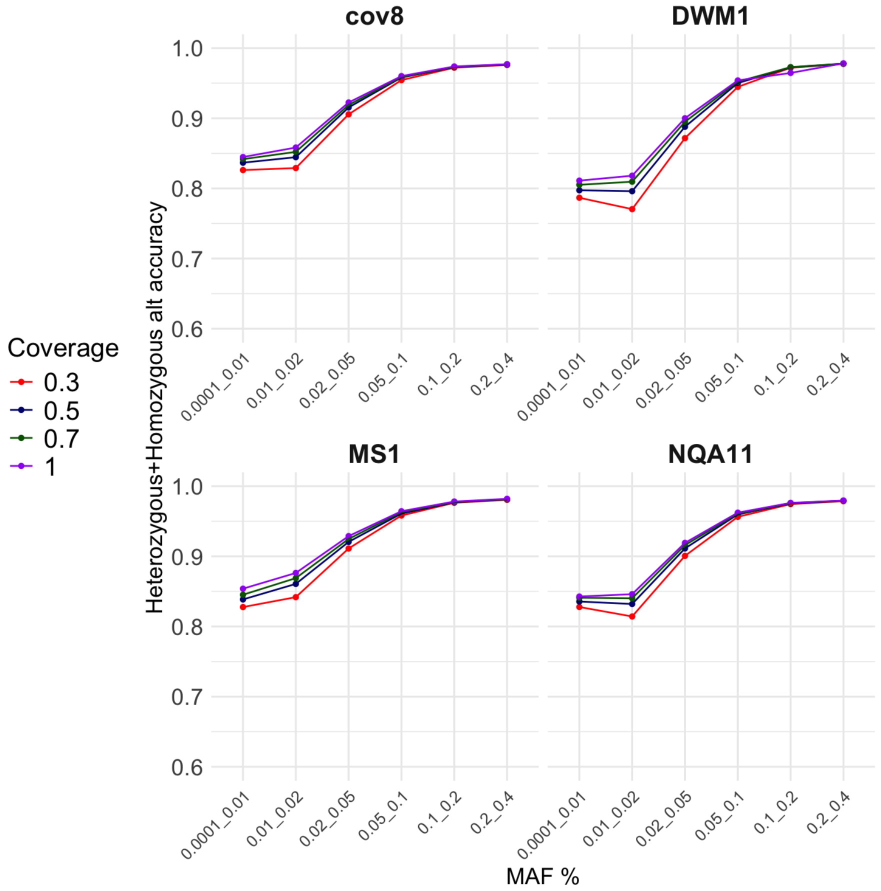


Figure S1: Accuracy rate of imputation of homozygous alternative and heterozygous sites (summed) for three ancient and two modern sheep downsampled to 0.3, 0.5, 0.7 and 1X genomic coverages (GP ≥0.99, INFO ≥0.99). Accuracy was investigated for different MAF bins (X-axis). Coverage is indicated by colour.

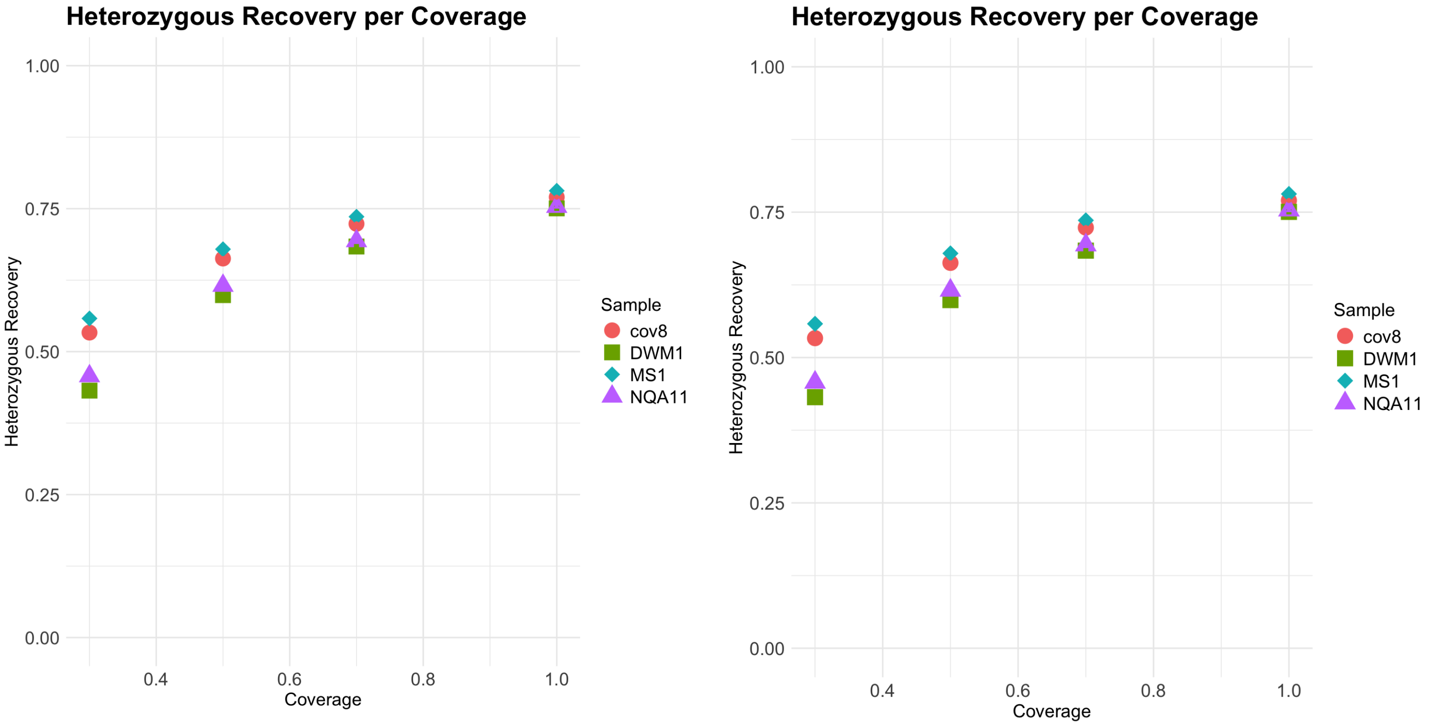


Figure S2: Recovery rates of imputed heterozygous and homozygous alternative sites for the test sheep at downsampled genomic coverages of 0.3, 0.5, 0.7 and 1X. Sample is denoted by shape and colour.

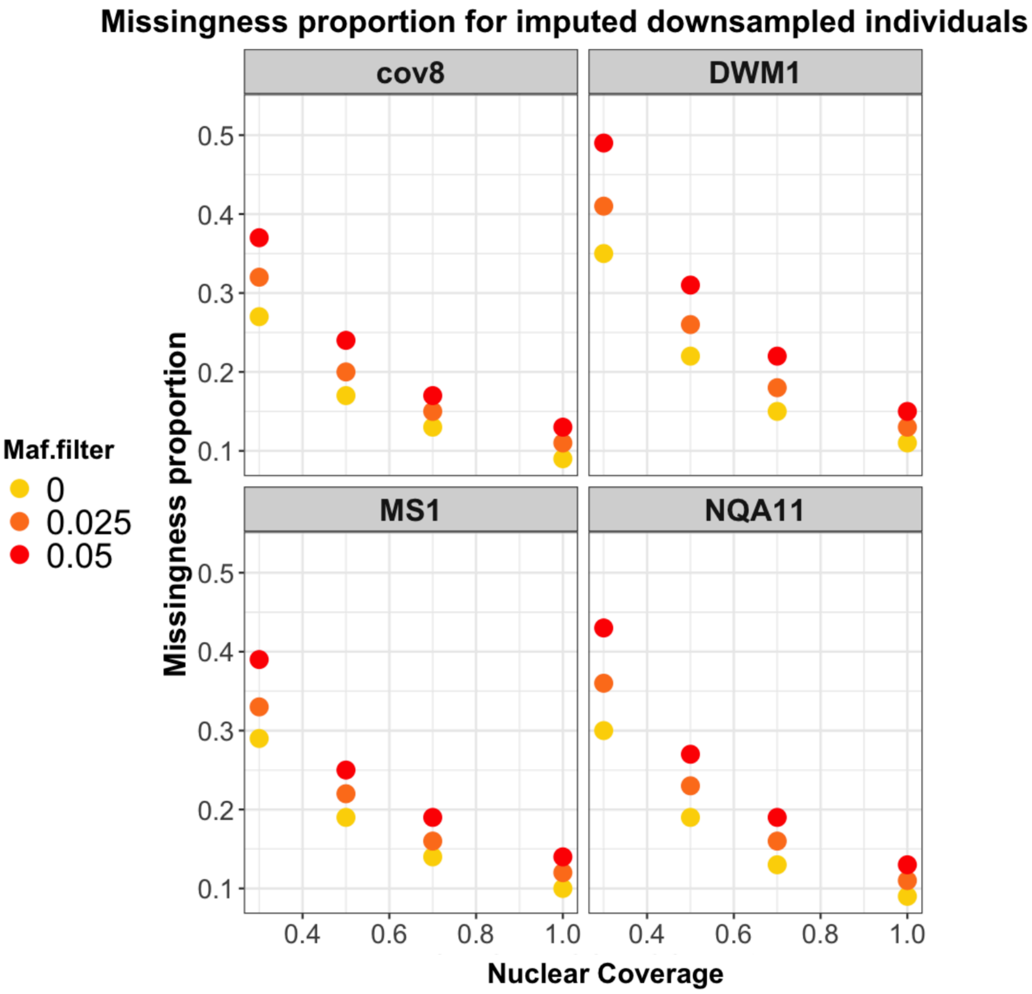


Figure S3: The proportion of missingness in each imputed sample at downsampled genomic coverages of 0.3, 0.5, 0.7 and 1X. This was measured for three different MAF thresholds (0, 0.025 and, 0.05) which are denoted by colour.

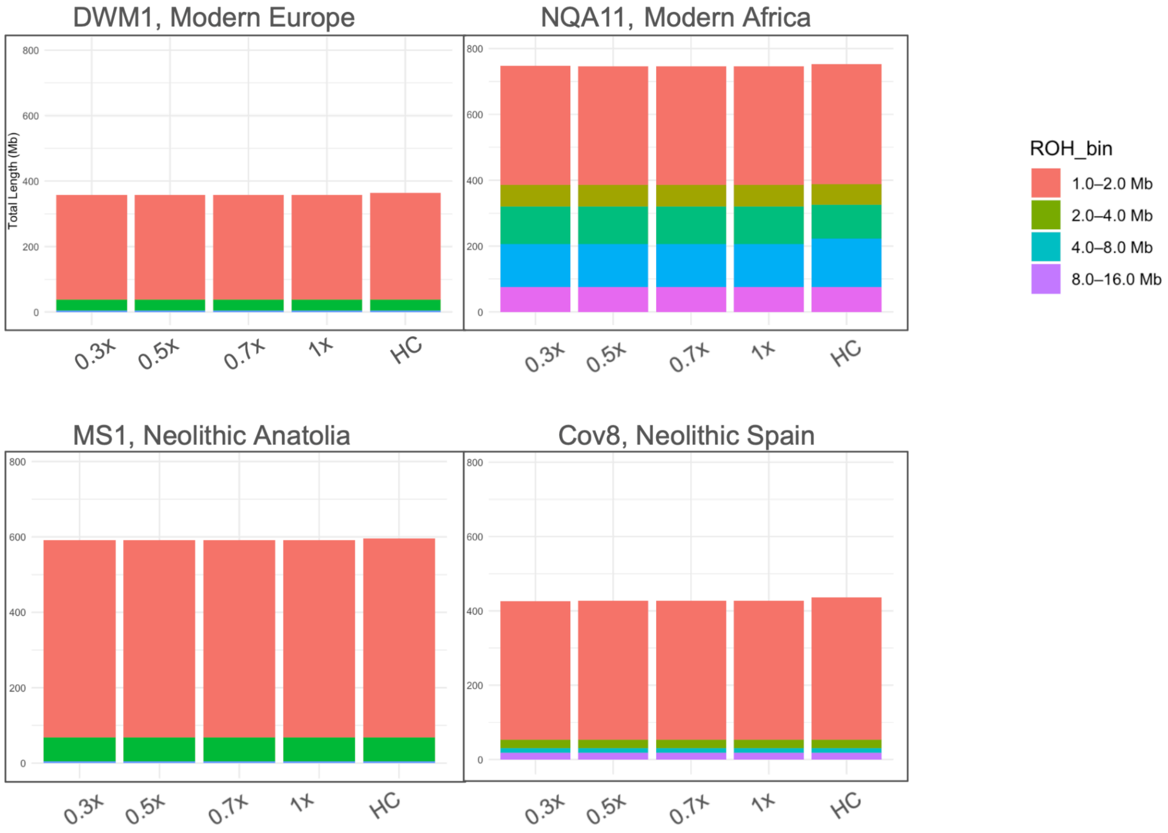


Figure S4: ROH estimates of downsampled imputed genomes and high-quality genotypes. The total ROH for each sample is divided into bins based on length (MAF >5%, no missingness, all sites).

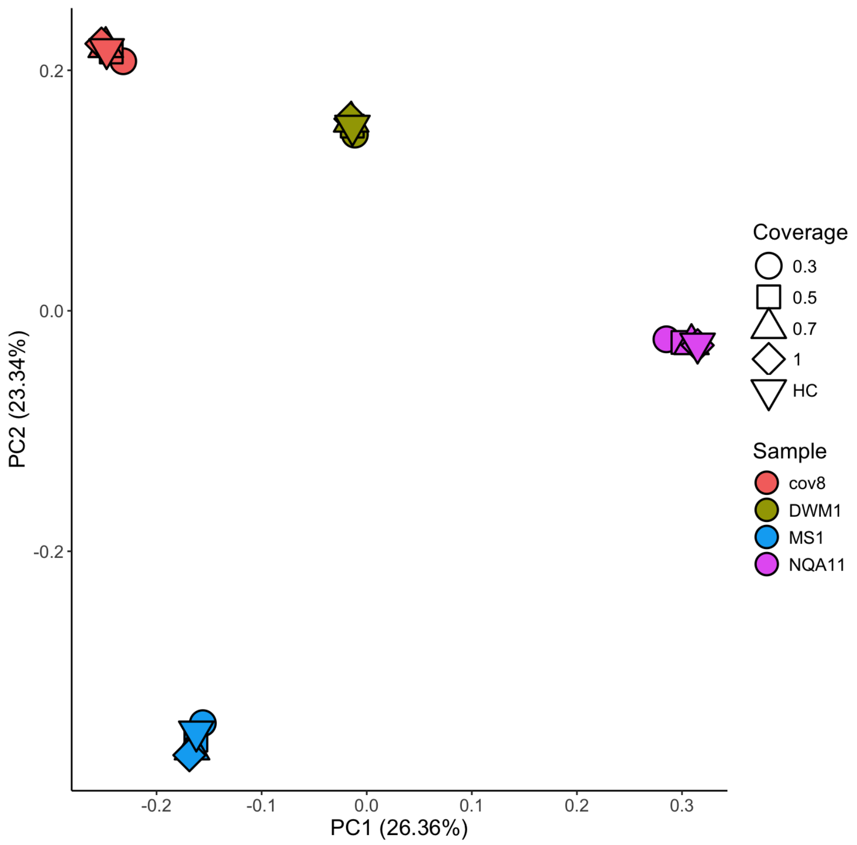


Figure S5. Principal component analysis of downsampled imputed genomes across different sequencing coverages. Samples cluster consistently across coverage levels, indicating that imputation results are robust to reductions in sequencing depth.

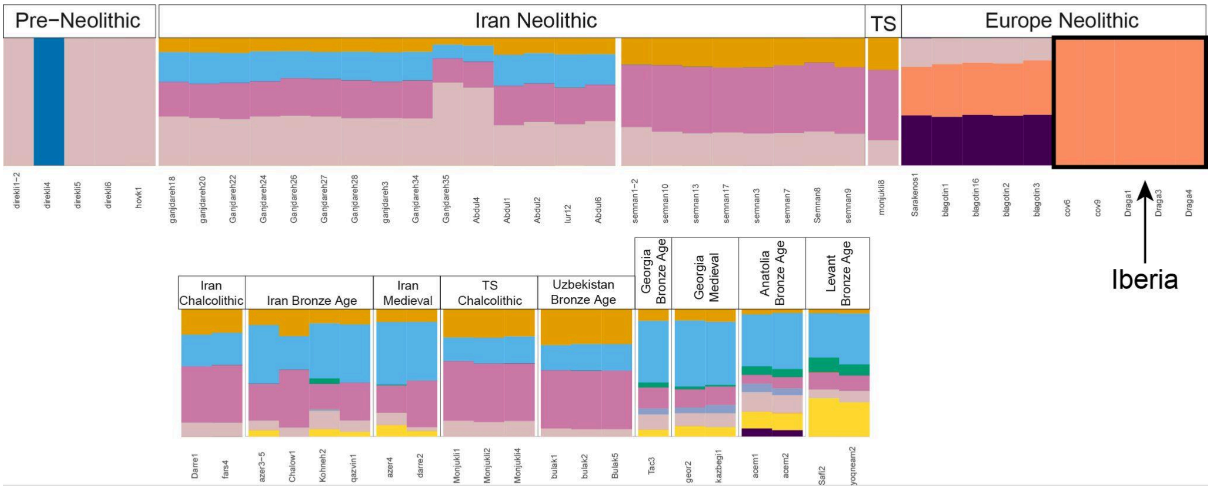


Fig. S6: ADMIXTURE profiles of all ancient goats, modern goats are shown in Table S11 with an optimal K of 14. ADMIXTURE was performed with pseudohaploid SNPS, filtered for transversion and a MAF of 5%, resulting in 2,420,382 SNPs.

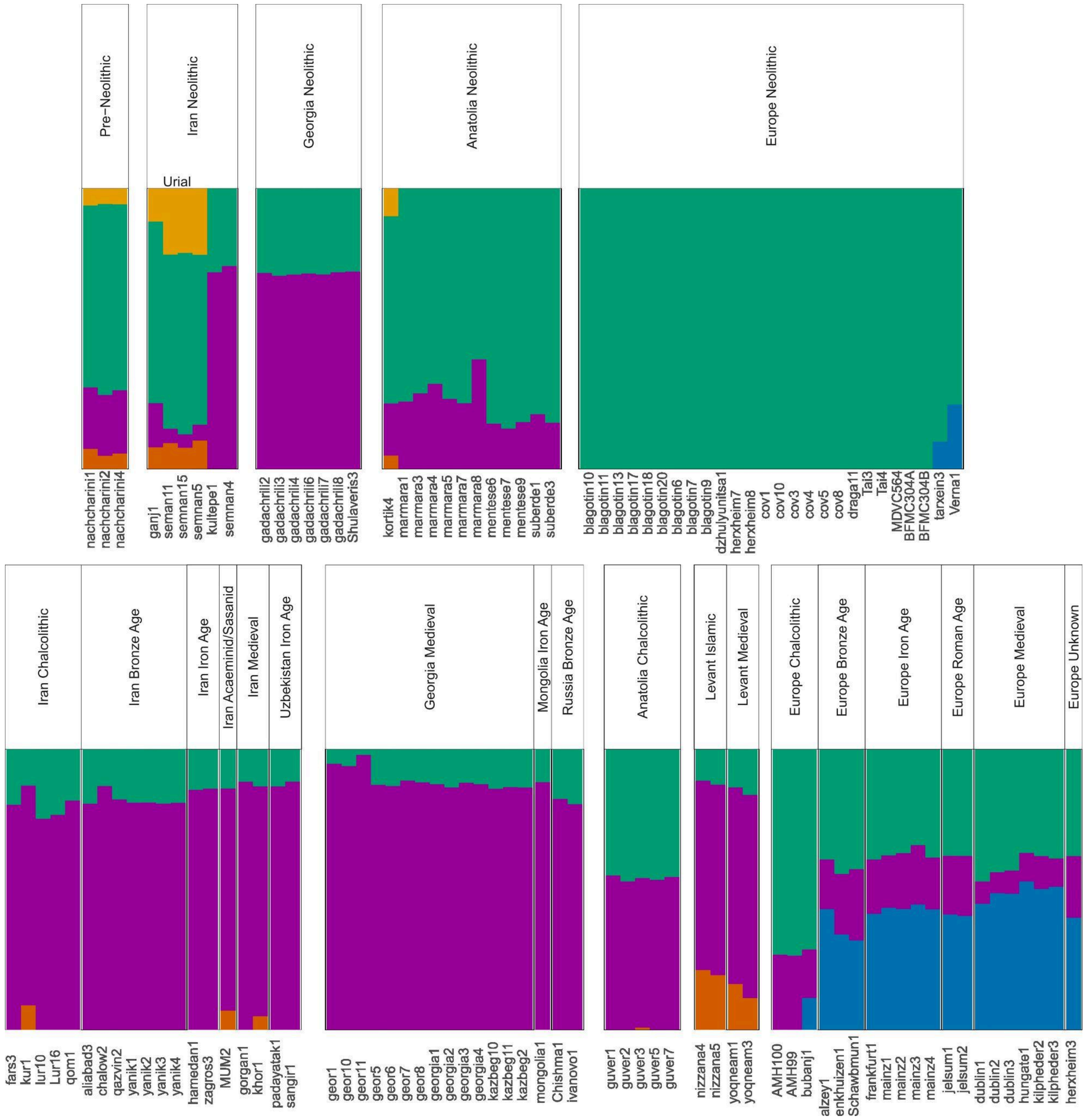

Figure S7: ADMIXTURE profiles of all ancient sheep, modern sheep are shown in Table S12 with an optimal K of 6. ADMIXTURE was performed with pseudohaploid SNPS, filtered for transversion and a MAF of 5%, resulting in 1,302,568 SNPs.

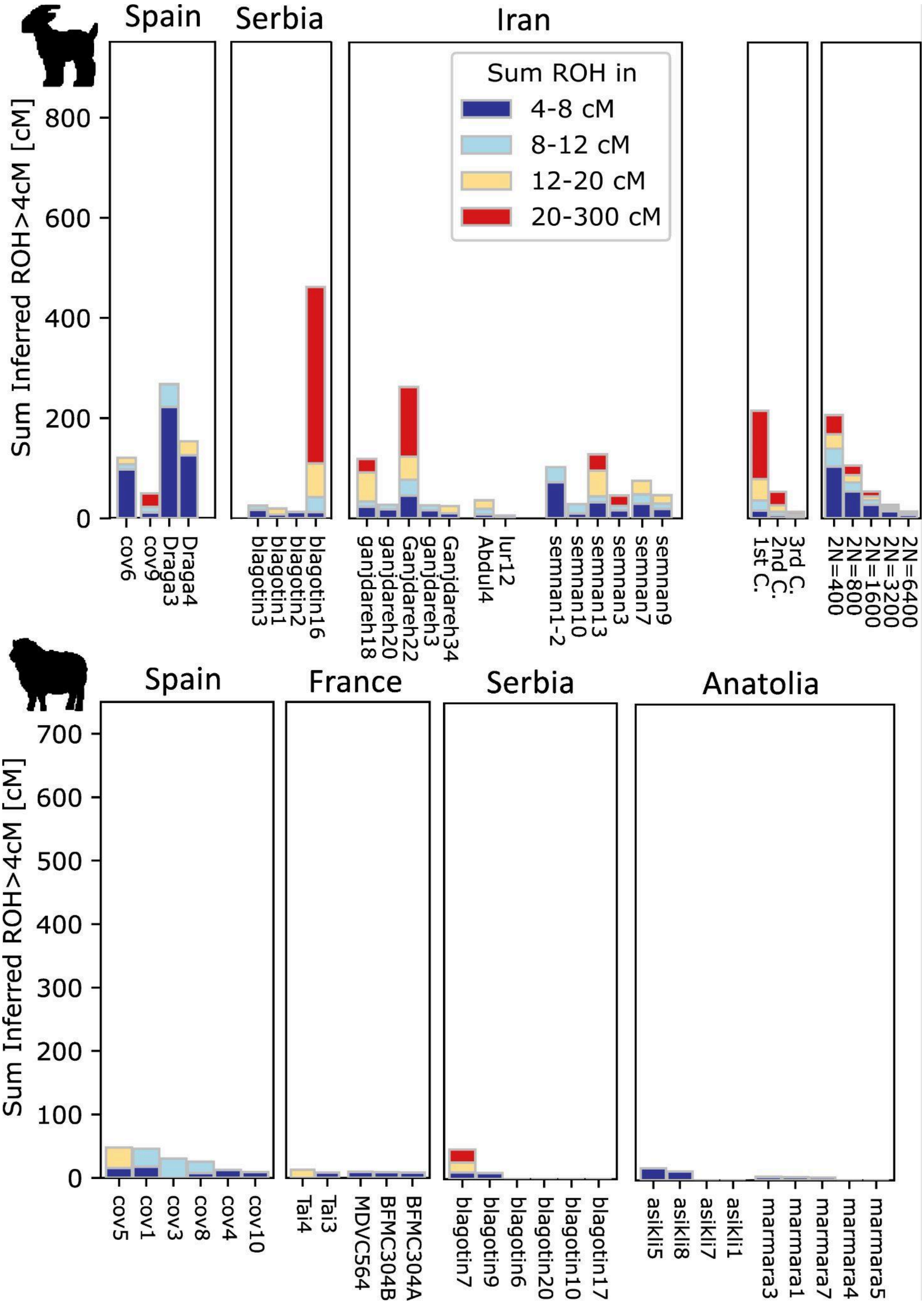


Figure S8: hapROH ROH profiles from Neolithic sheep and goats, using pseudohaploid variants. ROH was calculated with hapROH.

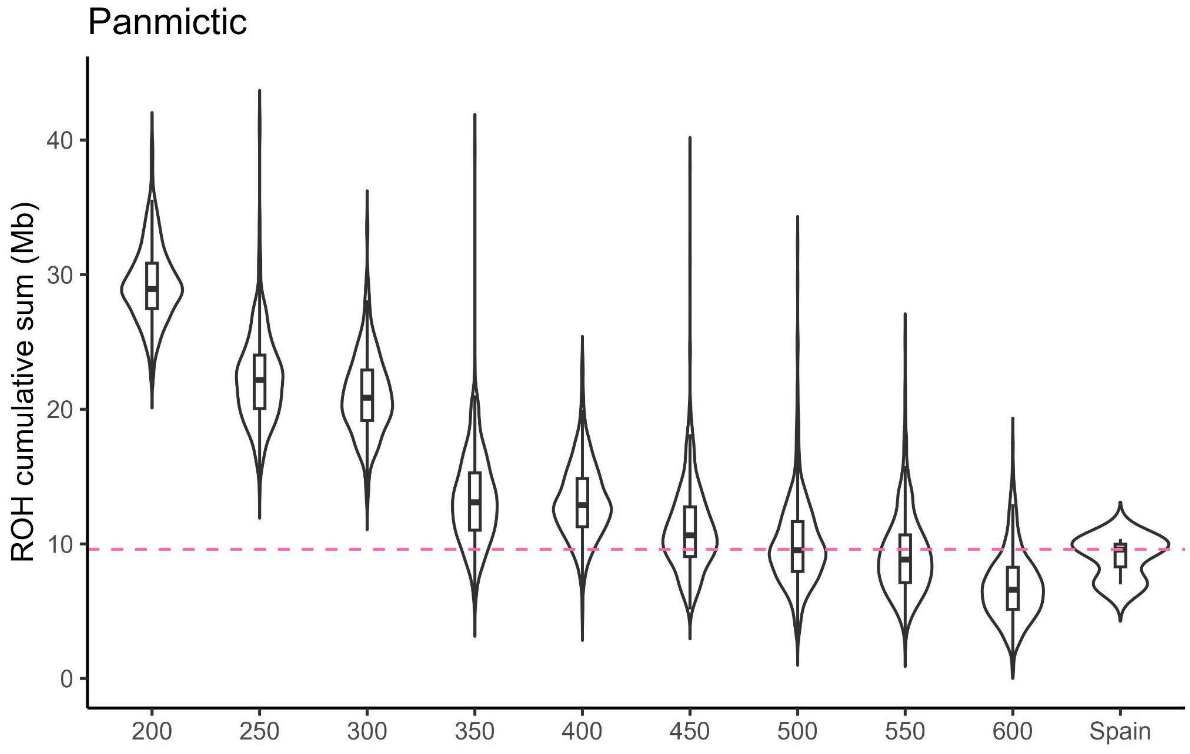


Figure S9: ROH profiles of a simulated panmictic population on chromosome 28. A simulation was performed mimicking goat demographic history (See SI methods) with a bottleneck of X, portrayed on the x-axis. ROH profiles were computed and total ROH length is shown on the y-axis. The pink dashed line is the average ROH profile of the empirical Iberian goat data.

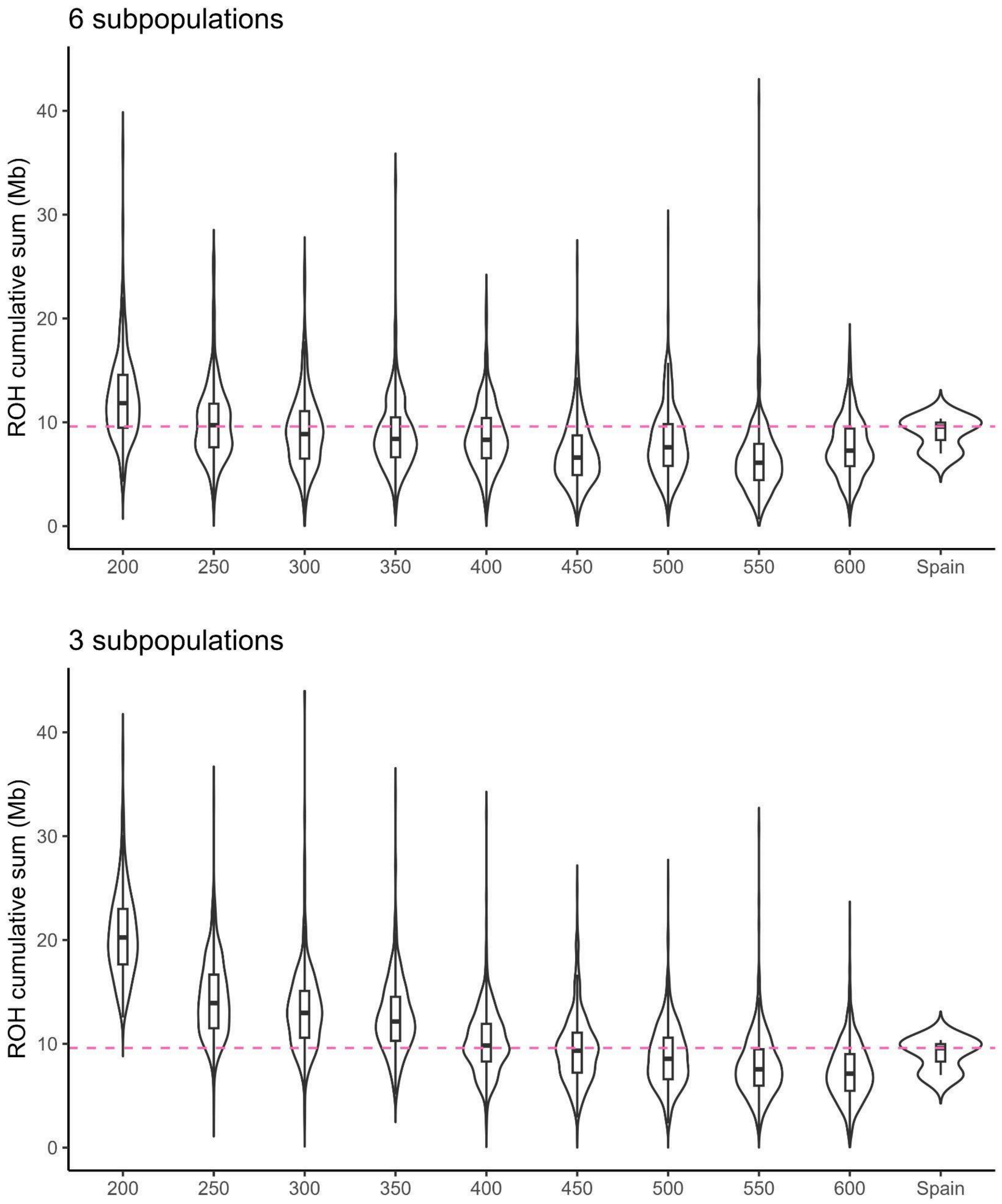


Figure S10: ROH profiles of a structured population (3 or 6 subpopulations) on chromosome 28. A simulation was performed mimicking goat demographic history (See SI methods) with a combined bottleneck of X, portrayed on the x-axis. ROH profiles were computed and total ROH length is shown on the y-axis. The pink dashed line is the average ROH profile of the empirical Iberian goat data.

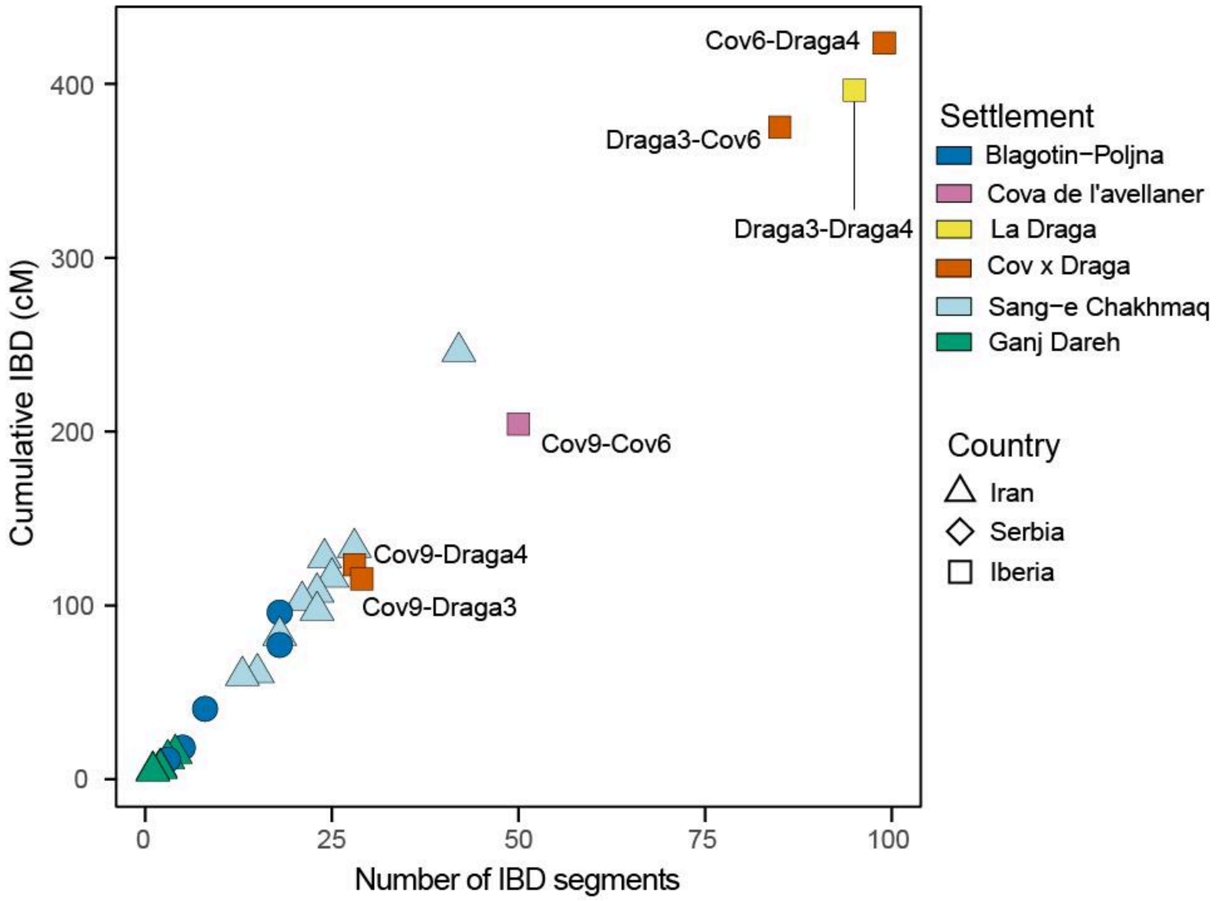


Figure S11: Pairwise IBD sharing within settlements for goats calculated with the strict pipeline (Table S7).

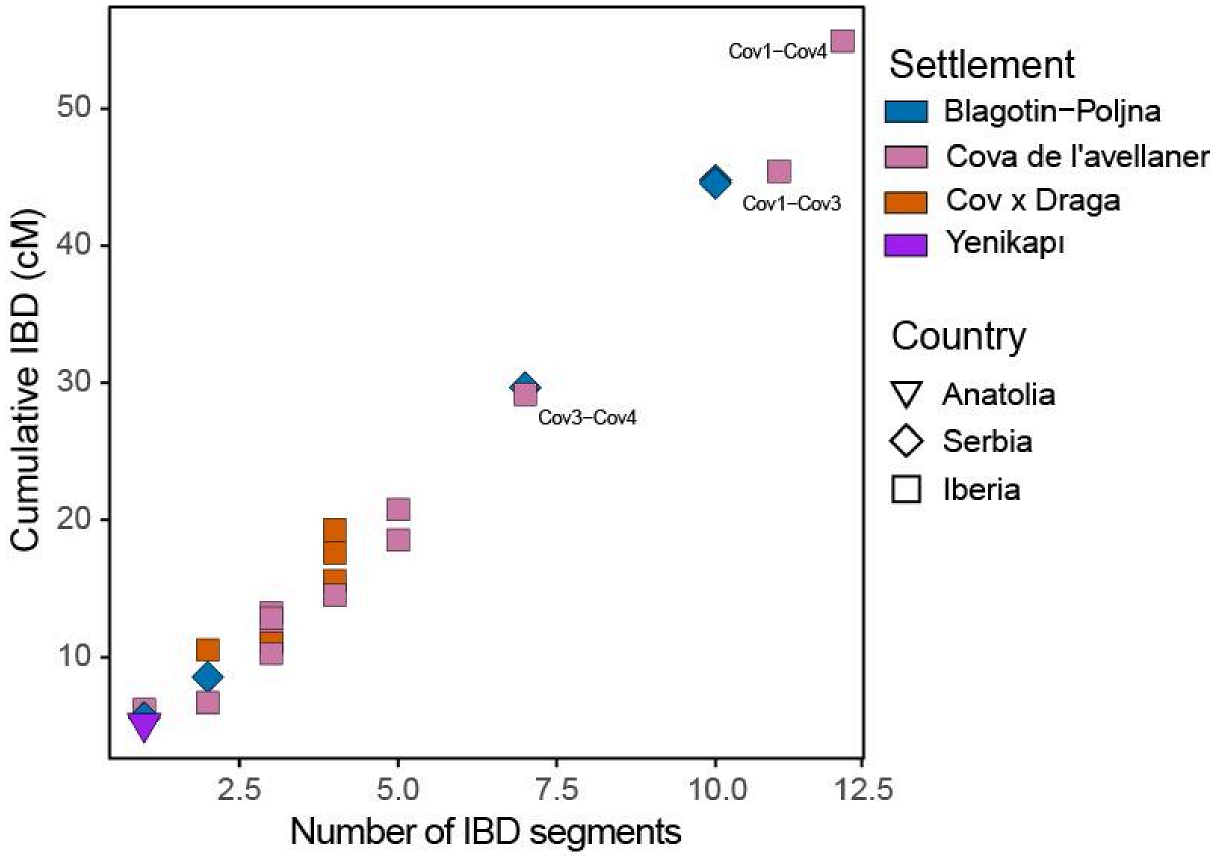


Figure S12: Pairwise IBD sharing within settlements for sheep calculated with the strict pipeline (Table S8).

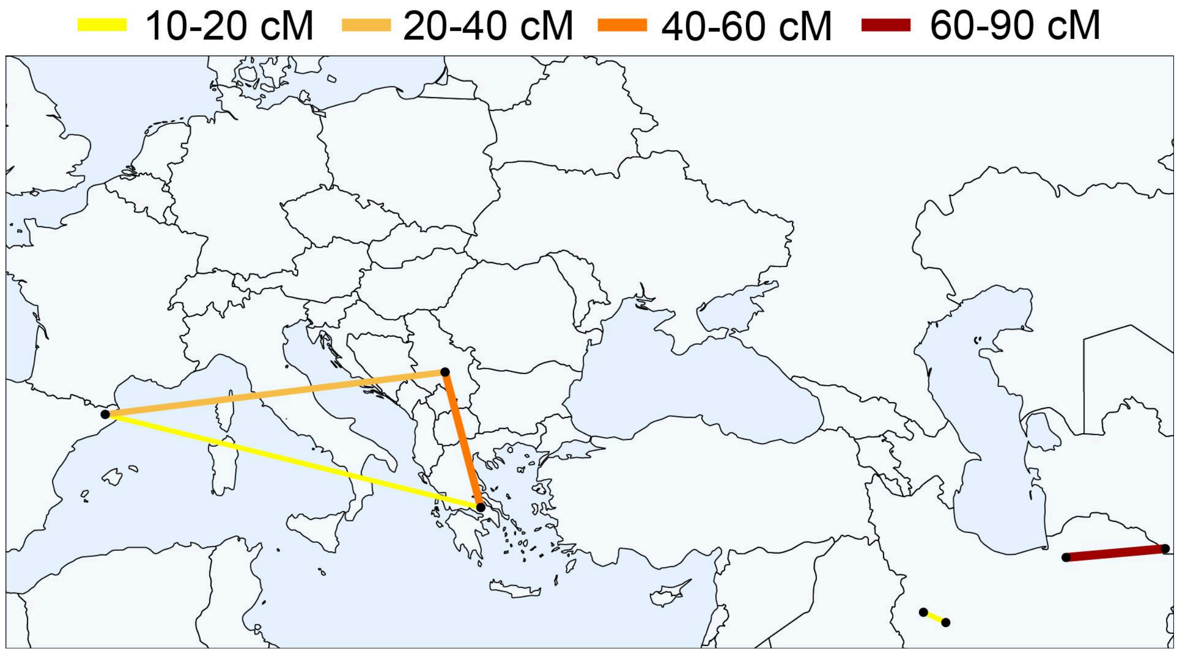


Figure S13: Between-settlement highest shared IBD (cM) for goat. Between-settlement IBD is represented by lines between-settlement pairs and does not imply direct cultural connectivity or dispersal patterns.

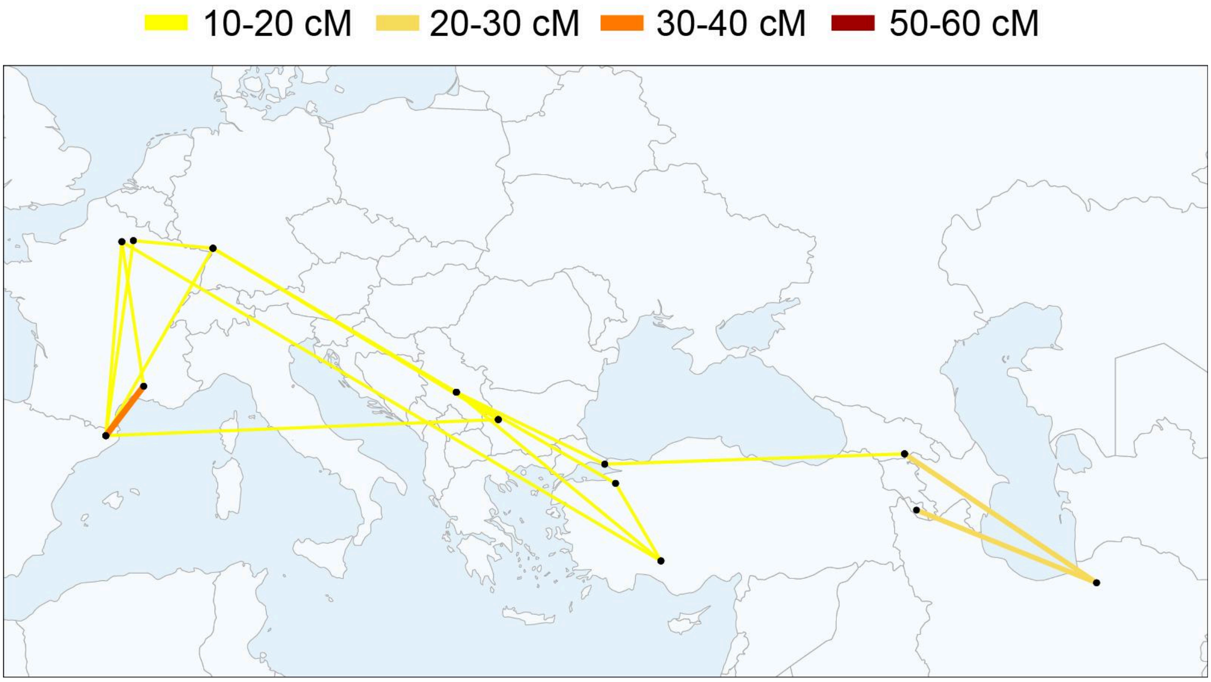


Figure S14: Between-settlement highest shared IBD (cM) for sheep. Between-settlement IBD is represented by lines between-settlement pairs and does not imply direct cultural connectivity or dispersal patterns.

Andreaki, V., J. A. Barceló, F. Antolín, et al. 2022. “Absolute Chronology at the Waterlogged Site of La Draga (lake Banyoles, Ne Iberia): Bayesian Chronological Models Integrating Tree-Ring Measurement, Radiocarbon Dates and Micro-Stratigraphical Data.” *Radiocarbon*, September 20, 1–42.

Andrews, Simon, and Others. 2010. “FastQC: A Quality Control Tool for High Throughput Sequence Data.” *Reference Source*.

Antolín, F., R. Buxó, S. Jacomet, V. Navarrete, and M. Saña. 2014. “An Integrated Perspective on Farming in the Early Neolithic Lakeshore Site of La Draga (Banyoles, Spain).” *Environmental Archaeology* 19: 241–255.

Bickhart, Derek M., Benjamin D. Rosen, Sergey Koren, et al. 2017. “Single-Molecule Sequencing and Chromatin Conformation Capture Enable de Novo Reference Assembly of the Domestic Goat Genome.” *Nature Genetics* 49 (4): 643–650.

Bosch, À., J. Chinchilla, and J. Tarrús, eds. 2011. *El Poblat Lacustre Del Neolític Antic de La Draga. Excavacions 2000--2005*. Vol. 9. Monografies Del CASC. Museu d’Arqueologia de Catalunya.

Bosch, A., and J. Tarrús. 1990. “La Cova Sepulcral Del Neolític Antic de l’Avellaner, Cogolls, Les Planes d'Hostoles (La Garrotxa).” In *Serie Monografica*, vol. 11, 11. Centre d’Investigacions Arqueologiques de Girona.

Bréhard, Stéphanie, and Jean-Denis Vigne. 2022. “Exploitation de La Faune Mammalienne et Caractérisation Des Occupations Du Néolithique Ancien et Moyen Du Site Du Ta"i.” In *Les Premières Sociétés Agropastorales Du Languedoc Méditerranéen. Le Ta"i (Remoulins -- Gard)*, edited by Claire Manen. Archives d’Écologie Préhistorique.

Briggs, Adrian W., Udo Stenzel, Matthias Meyer, Johannes Krause, Martin Kircher, and Svante Pääbo. 2010. “Removal of Deaminated Cytosines and Detection of in Vivo Methylation in Ancient DNA.” *Nucleic Acids Research* 38 (6): e87.

Browning, Brian L., and Sharon R. Browning. 2013. “Improving the Accuracy and Efficiency of Identity-by-Descent Detection in Population Data.” *Genetics* 194 (2): 459–471.

Cassidy, Lara M., Matthew D. Teasdale, Seán Carolan, et al. 2017. “Capturing Goats: Documenting Two Hundred Years of Mitochondrial DNA Diversity among Goat Populations from Britain and Ireland.” *Biology Letters* 13 (3). https://doi.org/10.1098/rsbl.2016.0876.

Chang, Christopher C., Carson C. Chow, Laurent Cam Tellier, Shashaank Vattikuti, Shaun M. Purcell, and James J. Lee. 2015. “Second-Generation PLINK: Rising to the Challenge of Larger and Richer Datasets.” *GigaScience* 4 (1): 7.

Colli, Licia, Hovirag Lancioni, Irene Cardinali, et al. 2015. “Whole Mitochondrial Genomes Unveil the Impact of Domestication on Goat Matrilineal Variability.” *BMC Genomics* 16 (1): 1115.

Daly, Kevin G., Pierpaolo Maisano Delser, Victoria E. Mullin, et al. 2018. “Ancient Goat Genomes Reveal Mosaic Domestication in the Fertile Crescent.” *Science* 361 (6397): 85–88.

Daly, Kevin G., Valeria Mattiangeli, Andrew J. Hare, et al. 2021. “Herded and Hunted Goat Genomes from the Dawn of Domestication in the Zagros Mountains.” *Proceedings of the*

*National Academy of Sciences of the United States of America* 118 (25). https://doi.org/10.1073/pnas.2100901118.

Damgaard, Peter B., Ashot Margaryan, Hannes Schroeder, Ludovic Orlando, Eske Willerslev, and Morten E. Allentoft. 2015. "Improving Access to Endogenous DNA in Ancient Bones and Teeth." *Scientific Reports* 5 (1): 11184.

Danecek, Petr, Adam Auton, Goncalo Abecasis, et al. 2011. "The Variant Call Format and VCFtools." *Bioinformatics (Oxford, England)* 27 (15): 2156–2158.

Danecek, Petr, James K. Bonfield, Jennifer Liddle, et al. 2021. "Twelve Years of SAMtools and BCFtools." *GigaScience* 10 (2). https://doi.org/10.1093/gigascience/giab008.

Davenport, Kimberly M., Derek M. Bickhart, Kim Worley, et al. 2022. "An Improved Ovine Reference Genome Assembly to Facilitate in-Depth Functional Annotation of the Sheep Genome." *GigaScience* 11 (February): giab096.

Denoyelle, Laure, Estelle Talouarn, Philippe Bardou, et al. 2021. "VarGoats Project: A Dataset of 1159 Whole-Genome Sequences to Dissect Capra Hircus Global Diversity." *Genetics, Selection, Evolution: GSE* 53 (1): 86.

Edgar, Robert C. 2004. "MUSCLE: Multiple Sequence Alignment with High Accuracy and High Throughput." *Nucleic Acids Research* 32 (5): 1792–1797.

Eggertsson, Hannes P., Hakon Jonsson, Snaedis Kristmundsdottir, et al. 2017a. "Graphtyper Enables Population-Scale Genotyping Using Pangenome Graphs." *Nature Genetics* 49 (11): 1654–1660.

Eggertsson, Hannes P., Hakon Jonsson, Snaedis Kristmundsdottir, et al. 2017b. "Graphtyper Enables Population-Scale Genotyping Using Pangenome Graphs." *Nature Genetics* 49 (11): 1654–1660.

Erven, Jolijn A. M., Alice Etourneau, Marjan Mashkour, et al. 2025. "Inferring Domestic Goat Demographic History through Ancient Genome Imputation." In *bioRxiv*. April 21. https://doi.org/10.1101/2025.04.18.649576.

Erven, Jolijn A. M., Amelie Scheu, Marta Pereira Verdugo, et al. 2024. "A High-Coverage Mesolithic Aurochs Genome and Effective Leveraging of Ancient Cattle Genomes Using Whole Genome Imputation." *Molecular Biology and Evolution* 41 (5). https://doi.org/10.1093/molbev/msae076.

Etourneau, Alice, Rachel Rupp, and Bertrand Servin. 2025. "Genome Landscape and Genetic Architecture of Recombination in Domestic Goats (Capra Hircus)." In *bioRxiv*. May 15. https://doi.org/10.1101/2025.05.15.654186.

Garcia-Reig, S. 2016. "Investigating Early Introduction of Sheep in the Iberian Peninsula: Reconstructing Seasonal Reproductive Patterns by Sequential Stable Isotope Analyses ( delta^18O) of Sheep Specimens from Cova de l'Avellaner (Early Neolithic – Catalonia)." Universitat Autònoma de Barcelona.

Gibaja, F., Berta Morell, Diego López-Onaindia, et al. 2018. "Nuevos Datos Cronológicos Sobre La Cueva Sepulcral Neolítica de l'Avellaner (Les Planes d'Hostoles, Girona)." *Munibe Antropologia-Arkeologia*, ahead of print, June 5. https://doi.org/10.21630/maa.2018.69.01.

Gouy, Manolo, Stéphane Guindon, and Olivier Gascuel. 2010. “SeaView Version 4: A Multiplatform Graphical User Interface for Sequence Alignment and Phylogenetic Tree Building.” *Molecular Biology and Evolution* 27 (2): 221–224.

Guindon, Stéphane, Jean-François Dufayard, Vincent Lefort, Maria Anisimova, Wim Hordijk, and Olivier Gascuel. 2010. “New Algorithms and Methods to Estimate Maximum-Likelihood Phylogenies: Assessing the Performance of PhyML 3.0.” *Systematic Biology* 59 (3): 307–321.

Haller, B. C., C. W. Nelson, M. F. Rodrigues, and P. W. Messer. 2025. “SimHumanity: Using SLiM 5.0 to Run Whole-Genome Simulations of Human Evolution.” In *bioRxivorg*. September 2. https://doi.org/10.1101/2025.09.01.673541.

Kaptan, Damla, Gözde Atağ, Kıvılcım Başak Vural, et al. 2024. “The Population History of Domestic Sheep Revealed by Paleogenomes.” *Molecular Biology and Evolution* 41 (10): msae158.

Katoh, Kazutaka, Kazuharu Misawa, Kei-Ichi Kuma, and Takashi Miyata. 2002. “MAFFT: A Novel Method for Rapid Multiple Sequence Alignment Based on Fast Fourier Transform.” *Nucleic Acids Research* 30 (14): 3059–3066.

Korneliussen, Thorfinn Sand, Anders Albrechtsen, and Rasmus Nielsen. 2014. “ANGSD: Analysis of Next Generation Sequencing Data.” *BMC Bioinformatics* 15 (November): 356.

Lacan, Marie, Christine Keyser, François-Xavier Ricaut, et al. 2011. “Ancient DNA Suggests the Leading Role Played by Men in the Neolithic Dissemination.” *Proceedings of the National Academy of Sciences of the United States of America* 108 (45): 18255–18259.

Li, Heng. 2011. “A Statistical Framework for SNP Calling, Mutation Discovery, Association Mapping and Population Genetical Parameter Estimation from Sequencing Data.” *Bioinformatics (Oxford, England)* 27 (21): 2987–2993.

Li, Heng, and Richard Durbin. 2009. “Fast and Accurate Short Read Alignment with Burrows-Wheeler Transform.” *Bioinformatics* 25 (14): 1754–1760.

Li, Heng, Bob Handsaker, Alec Wysoker, et al. 2009. “The Sequence Alignment/Map Format and SAMtools.” *Bioinformatics* 25 (16): 2078–2079.

Manen, Claire, ed. 2022. *Les Premières Sociétés Agropastorales Du Languedoc Méditerranéen. Le Taï (Remoulins -- Gard)*. Archives d’Écologie Préhistorique.

Meyer, Matthias, and Martin Kircher. 2010. “Illumina Sequencing Library Preparation for Highly Multiplexed Target Capture and Sequencing.” *Cold Spring Harbor Protocols* 2010 (6): db.prot5448.

Molina i Serramitjana, J. A. 1990. “Capítol V: Anàlisi de La Fauna.” In *La Cova Sepulcral Del Neolític Antic de l’Avellaner (Cogolls, Les Planes d’Hostoles, La Garrotxa)*, edited by À. Bosch i Lloret and J. Tarrús i Galter, vol. 11, 11. Sèrie Monogràfica de Girona. Centre d’Investigacions Arqueològiques de Girona.

Okonechnikov, Konstantin, Ana Conesa, and Fernando García-Alcalde. 2016. “Qualimap 2: Advanced Multi-Sample Quality Control for High-Throughput Sequencing Data.” *Bioinformatics (Oxford, England)* 32 (2): 292–294.

Palomo, Antoni, Raquel Piqué, Xavier Terradas, et al. 2014. “Prehistoric Occupation of Banyoles Lakeshore: Results of Recent Excavations at La Draga Site, Girona, Spain.” *Journal of Wetland Archaeology* 14 (1): 58–73.

Park, Stephen D. E., David A. Magee, Paul A. McGettigan, et al. 2015. “Genome Sequencing of the Extinct Eurasian Wild Aurochs, Bos Primigenius, Illuminates the Phylogeography and Evolution of Cattle.” *Genome Biology* 16 (October): 234.

Patterson, Nick, Priya Moorjani, Yontao Luo, et al. 2012. “Ancient Admixture in Human History.” *Genetics* 192 (3): 1065–1093.

Piqué, Raquel, Antoni Palomo, Xavier Terradas, et al. 2021. “Models of Neolithisation of Northeastern Iberian Peninsula: New Evidence of Human Occupations during the Sixth Millennium Cal BC.” *Open Archaeology* 7 (1): 671–689.

Quinlan, Aaron R., and Ira M. Hall. 2010. “BEDTools: A Flexible Suite of Utilities for Comparing Genomic Features.” *Bioinformatics* 26 (6): 841–842.

Ringbauer, Harald, John Novembre, and Matthias Steinrücken. 2021. “Parental Relatedness through Time Revealed by Runs of Homozygosity in Ancient DNA.” *Nature Communications* 12 (1): 5425.

Ripoll Miralda, J. 2025. “L’explotació I La Polivalència Ramadera Al Nord-Est de La Península Ibèrica: Canvis Productius Entre El VI I El IV Mil·lenni ANE.” Universitat Autònoma de Barcelona, PhD Programme in Prehistoric Archaeology.

Rubinacci, Simone, Robin J. Hofmeister, Bárbara Sousa da Mota, and Olivier Delaneau. 2023. “Imputation of Low-Coverage Sequencing Data from 150,119 UK Biobank Genomes.” *Nature Genetics* 55 (7): 1088–1090.

Rubinacci, Simone, Diogo M. Ribeiro, Robin J. Hofmeister, and Olivier Delaneau. 2021. “Efficient Phasing and Imputation of Low-Coverage Sequencing Data Using Large Reference Panels.” *Nature Genetics* 53 (1): 120–126.

Saña, M. 2011. “La Gestió Dels Recursos Animals.” In *El Poblat Lacustre Del Neolític Antic de La Draga. Excavacions 2000--2005*, edited by À. Bosch, J. Chinchilla, and J. Tarrús, vol. 9, 9. Monografies Del CASC. Museu d’Arqueologia de Catalunya.

Schubert, Mikkel, Stinus Lindgreen, and Ludovic Orlando. 2016. “AdapterRemoval v2: Rapid Adapter Trimming, Identification, and Read Merging.” *BMC Research Notes* 9 (February): 88.

Skoglund, Pontus, Swapan Mallick, Maria Cátira Bortolini, et al. 2015. “Genetic Evidence for Two Founding Populations of the Americas.” *Nature* 525 (7567): 104–108.

Stamatakis, Alexandros. 2014. “RAxML Version 8: A Tool for Phylogenetic Analysis and Post-Analysis of Large Phylogenies.” *Bioinformatics (Oxford, England)* 30 (9): 1312–1313.

Terradas, Xavier, Raquel Piqué, Antoni Palomo, et al. 2017. “Farming Practices in the Early Neolithic according to Agricultural Tools: Evidence from La Draga Site (northeastern Iberia).” In *Times of Neolithic Transition along the Western Mediterranean*. Springer International Publishing.

Wang, Chaolong, Xiaowei Zhan, Liming Liang, Gonçalo R. Abecasis, and Xihong Lin. 2015. “Improved Ancestry Estimation for Both Genotyping and Sequencing Data Using

Projection Procrustes Analysis and Genotype Imputation." *American Journal of Human Genetics* 96 (6): 926–937.

Wingett, Steven W., and Simon Andrews. 2018. "FastQ Screen: A Tool for Multi-Genome Mapping and Quality Control." *F1000Research* 7 (August): 1338.

Bochenski, Z.M., Tomek, T., Wertz, K., Kaczanowska M., Kozłowski J.K., Sampson A. 2018. Who ate the birds: the taphonomy of Sarakenos Cave, Greece. *Archaeol Anthropol Sci* 10, 1603–1615, https://doi.org/10.1007/s12520-017-0488-3

Sampson A. (ed). 2008. The Sarakenos Cave at Akrephnion, Beotia, Greece, vol. I: the Neolithic and the Bronze Age. University of the Aegean—Polish Academy of Arts and Science, Athens

Sampson A., Kozłowski J.K., Kaczanowska M., Budek A., Nadachowski A., Tomek T.,Miękina
M. 2009, Sarakenos Cave in Boeotia, from Palaeolithic to the Early Bronze Age, Eurasian prehistory 6(1-2), 199-231.

Wilczyński J., Tomek T., Nadachowski A., Miękina B., Rzebik-Kowalska B., Peresviet-Soltan A., Stworzewicz E., Szyndlar Z., Marciszak A., Lõugas L. 2016a. Sarakenos Cave (Greece, Beotia) - faunal record and environmental changes during Pleistocene and Holocene, In: M. Kaczanowska, J.K. Kozłowski and A. Sampson (eds), The Sarakenos Cave at Akraephnion, Boeotia, Greece, vol.II. The Early Neolithic, the Mesolithic and the Final Palaeolithic, Polska Akademia Umiejetności, Krakow, pp. 63-80.

Wilczyński J., Tomek T., Pryor A. 2016b. Sarakenos Cave (Greece, Beotia) – Archaeozoological record, In: M. Kaczanowska, J.K. Kozłowski and A. Sampson (eds), The Sarakenos Cave at Akraephnion, Boeotia, Greece, vol.II. The Early Neolithic, the Mesolithic and the Final Palaeolithic, Polska Akademia Umiejetności, Krakow, pp. 81-90.
